# Evolution end classification of *tfd* gene clusters mediating bacterial degradation of 2,4-dichlorophenoxyacetic acid (2,4-D)

**DOI:** 10.1101/2023.07.27.550777

**Authors:** Timur Iasakov

## Abstract

The *tfd* (*tfd*_I_ and *tfd*_II_) are gene clusters originally discovered in plasmid pJP4 which is involved in the bacterial degradation of 2,4-dichlorophenoxyacetic acid (2,4-D) via the ortho-cleavage pathway of chlorinated catechols. They share this activity, with respect to substituted catechols, with clusters *tcb* and *clc*. Although great efforts have been devoted over nearly forty years to exploring the structural diversity of these clusters, their evolution has been poorly resolved to date, and their classification is clearly obsolete. Employing comparative genomic and phylogenetic approaches revealed that all *tfd* clusters can be classified as one of four different types. The following four-type classification and new nomenclature are proposed: *tfd*_I_, *tfd*_II_, *tfd*_III_ and *tfd*_IV(A,B,C)_. Horizontal gene transfer between *Burkholderiales* and *Sphingomonadales* provided phenomenal linkage between *tfd*_I_, *tfd*_II_, *tfd*_III_ and *tfd*_IV_ type clusters and their mosaic nature. It is hypothesized that the evolution of *tfd* gene clusters proceeded within first (*tcb*, *clc* and *tfd*_I_), second (*tfd*_II_ and *tfd*_III_) and third (*tfd*_IV(A,B,C)_) evolutionary lineages in each of which the genes were clustered in specific combinations. Their clusterization has been discussed through the prism of hot spots and driving forces of various models, theories, and hypotheses of cluster and operon formation. Two hypotheses about series of gene deletions and displacements have also been proposed to explain the structural variations across members of clusters *tfd*_II_ and *tfd*_III_, respectively. Taking everything into account, these findings reconstructing the phylogeny of *tfd* clusters, have delineated their evolutionary trajectories, and allowed the contribution of various evolutionary processes to be assessed.

## Introduction

Currently *tfd* gene clusters are model objects for studying the microbial acquisition of xenobiotic degradation capacity. They encode for enzymes involved in the degradation of 2,4-dichlorophenoxyacetic acid (2,4-D) which has human health and ecological risks (Peterson et al., 2016; ATSDR, 2020), but is still used worldwide as an herbicide in agriculture (Islam et al., 2018). Continued exploration of microbial 2,4-D degradation in various environments in China (Xia et al., 2017; Xiang et al., 2020), Brazil (Brucha et al., 2021), Vietnam (Nguyen et al., 2021a; Nguyen et al., 2021b), Russia (Zharikova et al., 2021) and Japan (Hayashi et al., 2021) suggests that this problem is still in the spotlight.

Previously, under the name “TFD”, conjugative plasmids (TFD plasmids) containing genes of the 2,4-D and 4-chloro-2-methylphenoxyacetic acid (MCPA) degradation pathways were described (Don, Pemberton, 1981). At the moment, the abbreviation *tfd* designates two clusters of genes involved in the degradation of 2,4-D, which are located on the pJP4 plasmid of strain *Cupriavidus pinatubonensis* JMP134 (previously identified as *Ralstonia eutropha*, *Alcaligenes eutrophus*, *Waustersia eutropha* and *Cupriavidus necator*). Each of these clusters, *tfdB*_I_*F*_I_*E*_I_*D*_I_*C*_I_*T* (designated as *tfd*-I/*tfd*_I_/*tfd-I*/*tfd_I_*) and *tfdKB*_II_*F*_II_*E*_II_*C*_II_*D*_II_*R* (designated as *tfd*-II/*tfd*_II_/*tfd-II*/*tfd_II_*), encodes a core set of genes for the ortho-cleavage pathway of chlorocatechol. In the case of cluster *tfd*_II_, this core set of genes is extended by the *tfdA* and *tfdK* genes (Don et al., 1985; Streber et al., 1987; Harker et al., 1989; Leveau et al., 1998; Kaphammer et al., 1990; Laemmli et al., 2000; Trefault et al., 2004 and other).

The reactions and genes controlling 2,4-D degradation are as follow: acetate side chain cleavage of 2,4-D by α-ketoglutarate-dependent 2,4-D dioxygenase (*tfdA*) into 2,4-dichlorophenol (2,4-DCP); hydroxylation reaction of 2,4-DCP by 2,4-DCP hydroxylase (*tfdB*) into 3,5-dichlorocatechol (3,5-DCC); ortho-cleavage of 3,5-dichlorocatechol by chlorocatechol 1,2-dioxygenase (*tfdC*) to form 2,4-dichloro-cis,cis-muconate (2,4-dichloromuconic acid); conversion of 2,4-dichloro-cis,cis-muconate to 2-chlorodienelactone catalyzed by chloromuconate cycloisomerase (*tfdD*); conversion of 2-chlorodienelactone to 2-chloromaleylacetate by chlorodienelactone hydrolase (*tfdE*); conversion of 2-chloromaleylacetate to 3-oxoadipate via maleylacetate by chloromaleylacetate reductase and maleylacetate reductase (*tfdF*), respectively, which is further funnelled to the tricarboxylic acid cycle (TCA) (Kumar et al., 2016). Functionally corresponding proteins of both clusters are complementary, since they have different efficiencies in terms of the performed reactions (Pérez-Pantoja et al., 2000; Laemmli et al., 2002; Plumeier et al., 2002).

An ortho-cleavage pathway of chlorinated catechols is a common catabolic function between *tfd* gene clusters on the one hand and *tcb* with *clc* gene clusters on the other hand. The *tcb* gene clusters are located on catabolic plasmid pP51 which contains two gene clusters, *tcbAB* and *tcbRCDEF*. The *tcbAB* controls the degradation of chlorinated benzenes to chlorinated catechols, while *tcbRCDEF* are directly responsible for the degradation of chlorinated catechols (van der Meer et al., 1991b; van der Meer et al., 1991c). The *clc* gene cluster (chlorocatechols – Clc) of plasmid pAC27 with a *clcRABDE* architecture controls the degradation of 3-chlorobenzoate (3-CB) via the ortho-cleavage pathway of chlorinated catechols (Frantz, Chakrabarty, 1987; Ghosal, You, 1989; Coco et al., 1993). It has been shown that functionally related proteins of *tfd*, *tcb* and *clc* gene clusters have high levels of similarity and identity between their amino acid and nucleotide sequences. Additionally, all clusters have shown strong DNA homology and similar organization (Ghosal et al., 1985; Ghosal, You, 1988; van der Meer et al., 1991 (a); Kasberg et al., 1995). Schlömann (1994) noted that phylogenetically, *tfd-I*, *tcb* and *clc* gene clusters diverged from a common ancestral pathway of chlorocatechols. But further phylogenetic analyses have indicated independent recruitment of diverse genes during assemblage of the *tfd* gene cluster (Vallaeys et al., 1999) and the probable independent evolution of the *tfdA* gene (Zharikova et al., 2018b).

In the past three decades, all explorations of *tfd* gene clusters, from a comparative genomic point of view, have been focused primarily on their tandem matching with the canonical gene structure of *tfd*_I_ и *tfd*_II_ gene clusters. A lot of both canonical and non-canonical clusters have been identified, amongst them: *tfdT-CDEF* (Liu et al., 2001), *tfdDRCEBKAF* (Poh et al., 2002), *tfdCDEF* and *tfdC*_II_*E*_II_*BKA* (*Delftia acidovorans* P4a), *tfdRCEBKA* (Vedler et al., 2004), *tfdC2E2F2* and *tfdDRFCE* (Thiel et al., 2005), *tfdAKBECR* (Sen et al. 2011), *tfdRDCEFBKA* (Kim et al., 2013), *tfdBCDEFKR* (Nielsen et al., 2013), *tfdFAKB*_I_*E*_I_*C*_I_*D*_I_*R*_I_ and *tfdC*_II_*E*_II_*B*_II_ (Sakai et al., 2014), *tfdFEDCT* (Ricker et al., 2016), *tfdB*_I_*F*_I_*E*_I_*D*_I_*C*_I_*T* and variations of the *tfdKB*_II_*FE*_II_*C*_II_*D*_II_*R* cluster with deletions (Nguyen et al., 2019), *tfdBFET*, *tfdAKBFECDR* and *tfdBFEDC* (Xiang et al., 2020), *tfdBFEDCS*, *tfdFE(tctC)DCS* (Nguyen et al., 2021a), *tfdBaFRDEC* (Hayashi et al., 2021), *tfdCDEF* (Yamamoto-Tamura et al., 2021), *tfdEICIFIRDI*,*K*,*BI*, *tfdDII*,*FIIEIICII*,*BII* (Zhang et al., 2023) and other cluster structures. Amongst the clusters related to *tcb* and *clc*, the following have been identified: *tetRtetCDEF* (Potrawfke et al., 2001), *mocpDCBAR* (Sen et al., 2011), *cbnR-ABCD* (Ogawa, Miyashita, 1999; Moriuchi et al., 2019), *clcRABCD* (Xiao et al., 2007), *clcRABCDE* (Jiang et al., 2009), *clcr1a1b1d1e1* (Miyazaki et al., 2015), *dccEDCBAR* (Li et al., 2021) and others. While the studies referred to above have been beneficial for gaining knowledge on a great variety of gene clusters structures controlling the chlorocatechol ortho-cleavage pathway (belonging to the *tfd*, *tcb* and *clc* gene clusters), nevertheless, their phylogeny and evolution are still poorly resolved, classification obsolete and the genes are still perhaps misannotated.

Comparative genomic and phylogenetic approaches were applied to fill these knowledge gaps. The findings allowed a new classification and nomenclature of these gene clusters, into *tfd*_I_, *tfd*_II_, *tfd*_III_ and *tfd*_IV (A, B, C)_ types, to be proposed. Additionally, the findings indicated that *tfd* are a unique family of mosaic clusters whose clusterization occurred in several evolutionary lineages with active recruitment of ancestral genes from *Burkholderiales* and *Sphingomonadales* through horizontal gene transfer. Possible paths for the clusterization of *tfd* genes within each lineage in the light of the various models of operon formation were discussed. These results constitute a further contribution to the understanding of bacterial genome organization and will be beneficial for the correct annotation of *tfd* clusters, as well as further studies of their diversity, propagation, and evolution.

## Materials and Methods

### Databases and data collection

All DNA and protein sequences, as well as additional information were obtained from both finished and unfinished genome sequencing records which are publicly available at NCBI resources (Sayers et al., 2023). The plasmids and genomes of 56 bacterial species were analyzed. Most of them belonged to the order *Burkholderiales* from Betaproteobacteria and a few belonged to Alpha- and Gammaproteobacteria.

### Gene annotation

Two ORF prediction softwares, the web version of the NCBI ORF finder and SnapGene Viewer v.4.2.1, were used. Then the predicted ORFs were annotated using NCBI blastp (Sayers et al., 2023) similarity searches against the UniProtKB/Swiss-Prot (swissprot) database.

### Comparative genomic analyses

The canonical gene clusters of the plasmids pJP4 (AY365053), pP51 (M57629) and pAC27 (M16964) were used as templates for NCBI blastn (Sayers et al., 2023) similarity searches against the nucleotide collection (nr/nt) and whole-genome shotgun contigs (wgs) databases. Adjacent to the clusters, genomic regions were searched and analyzed in order to determine synteny between clusters and flanked genes. For incomplete genome sequence projects in contig format, this strategy was not possible and gene cluster architecture was supposed based on nucleotide sequence identity.

Multiple sequence alignments of DNA and protein sequences of analyzed gene clusters were obtained using MAFFT at default parameter settings (Madeira et al., 2022). Values of identity and similarity between corresponding nucleotide and protein sequences of analyzed clusters were calculated using SIAS (http://imed.med.ucm.es/Tools/sias.html) with default parameters.

Comparative synteny analysis was performed by comparison between linearized maps and oriented according to the structure of gene clusters to depict gene arrangements, gene retention and orientation. Detailed synteny maps were visualized and blastn identity comparisons were generated using Easyfig v.2.2.2 (Sullivan et al., 2011).

The new classification scheme for *tfd* gene clusters into four types – I, II, III and IV (A, B and C) and nomenclature were proposed. An updated nomenclature of *tfd* gene clusters was formulated based on the following syntax: the three italicized lower case letters followed by Roman numerals as a subscript refer to the number of the cluster type. For example, *tfd*_I_*BFEDCT* (in short, *tfd*_I_), *tfd*_II_*AKBFECDR* (in short, *tfd*_II_), *tfd*_III_*FAKBECDR* (in short, *tfd*_III_), *tfd*_IVA_*ECFRD*,*B*/*tfd*_IVB_*CEF*,*D*/*tfd*_IVC_*CEDRF*,*B* (in short, *tfd*_IVA_, *tfd*_IVB_, *tfd*_IVC_). It is worth noting that there is no need to designate each gene as belonging to a certain type in the cluster structure (for example *tfdB*_I_*F*_I_*E*_I_*D*_I_*C*_I_*T*_I_), since *tfd* clusters do not formed hybrid clusters among themselves.

For clusters *tcb* and *clc*, a reverse spelling of the gene order, *tcbFEDCR* and *clcEDBAR*, was proposed, to correspond to the order of genes in other types of ortho-pathway clusters (*tfd*_I_, *tfd*_II_ and *tfd*_III_).

### Alignments and phylogenetic analyses

Multiple sequence alignments for all trees were performed with MAFFT using default parameter settings (Madeira et al., 2022). The evolutionary history was inferred by using the Maximum Likelihood method based on the JTT matrix-based model (Jones et al., 1992). Initial tree(s) for the heuristic search were obtained automatically by applying Neighbor-Join and BioNJ algorithms to a matrix of pairwise distances estimated using a JTT model, and then selecting the topology with a superior log likelihood value. Evolutionary analyses were conducted in MEGA7 (Kumar et al., 2016) with 1000 bootstrap replicates.

## Results

### Comparative genomics of *tfd*, *tcb* and *clc* gene clusters

The *tfd*, *tcb* и *clc* gene clusters were identified from publicly available genomes and plasmids by: i) using canonical gene clusters of the plasmids pJP4 (AY365053), pP51 (M57629) and pAC27 (M16964) as NCBI BLAST query sequences; ii) the analysis of published articles. Clusters searched for on 01/10/2023 which had two or more gene overlap matches were taken into account in analysis. In total, the above-mentioned search methods resulted in the identification of eighty-five *tfd*, *tcb* and *clc* genes clusters with different genetic structures.

### The *tfd*_I_ gene cluster

A blastn-search revealed seventeen (excluding canonical *tfd*_I_) gene clusters which had an over 90% match with the *tfdB*_I_*F*_I_*E*_I_*D*_I_*C*_I_*T* structure of *tfd*_I_ gene cluster. They were designated as *tfdT1T2CD* (unpublished), *tfdCDEF* or *tfdT-CDEF* (Liu et al., 2001; Yamamoto-Tamura et al., 2021), *tfdCDEF* (Morimoto, Fujii, 2009), *tfdFEDCT* (Ricker et al., 2016), *tfdB*_I_*F*_I_*E*_I_*D*_I_*C*_I_*T* (Nguyen et al., 2019), *tfdBFEDC*, *tfdBFET* (Xiang et al., 2020), *tfdBFEDCS* (Nguyen et al., 2021a) or not annotated (CP016005, CP026110) (Table 1). Fourteen of those seventeen *tfd*_I_ gene clusters were located on plasmids in species belonging to the *Burkholderiaceae* family. Amongst these plasmids were: pJP4 (AY365053), plasmid5 (=unnamed5 =plas5) (CP038640), a number of pPO line plasmid contigs (pPO1, 2, 3, 4, 7, 10, 16, 26, 27), pEMT1 (CP026110), pNK84 (AB050198) and pkk4 (CP016005). The three clusters were identified in genomes *Bordetella petrii* BT1 9.2 (WMBV01000066), uncultured bacterium (AB478351) and chromosome 1 of *Paraburkholderia phytofirmans* OLGA172 (CP014578). Both identified species belonged to the *Alcaligenaceae* and *Burkholderiaceae* families of the order *Burkholderiales*, respectively (table 1).

**Table 1.**
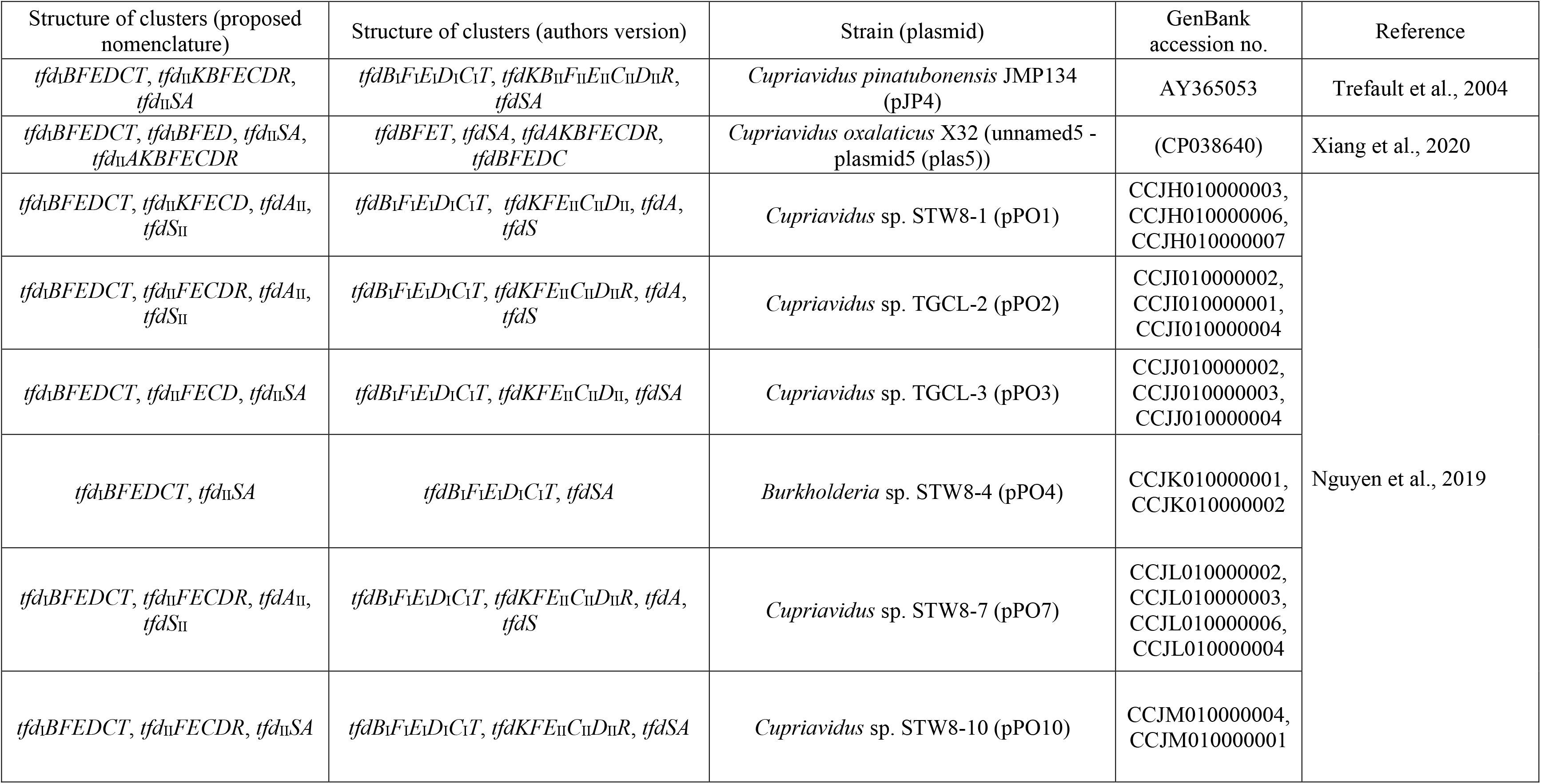
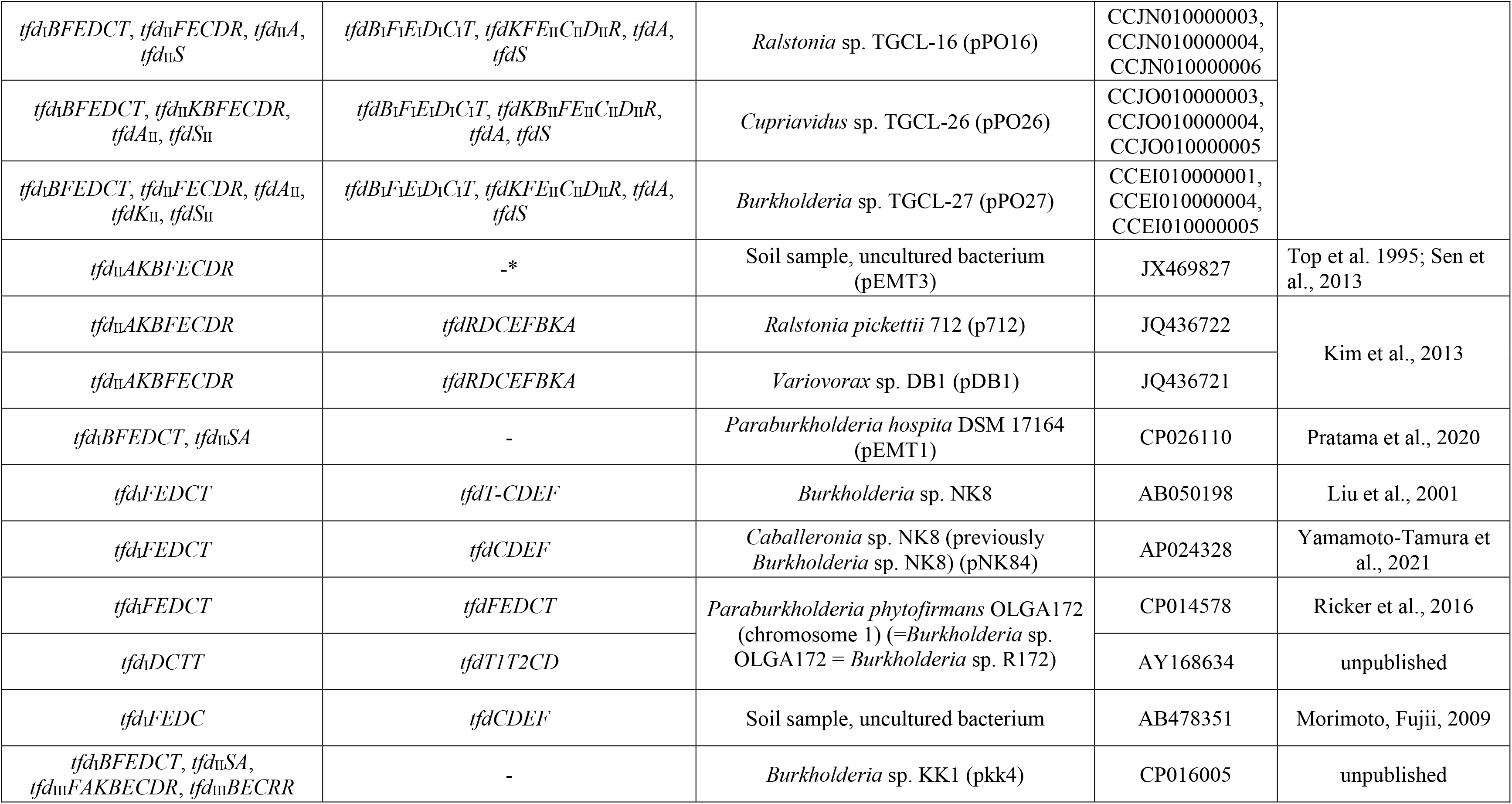
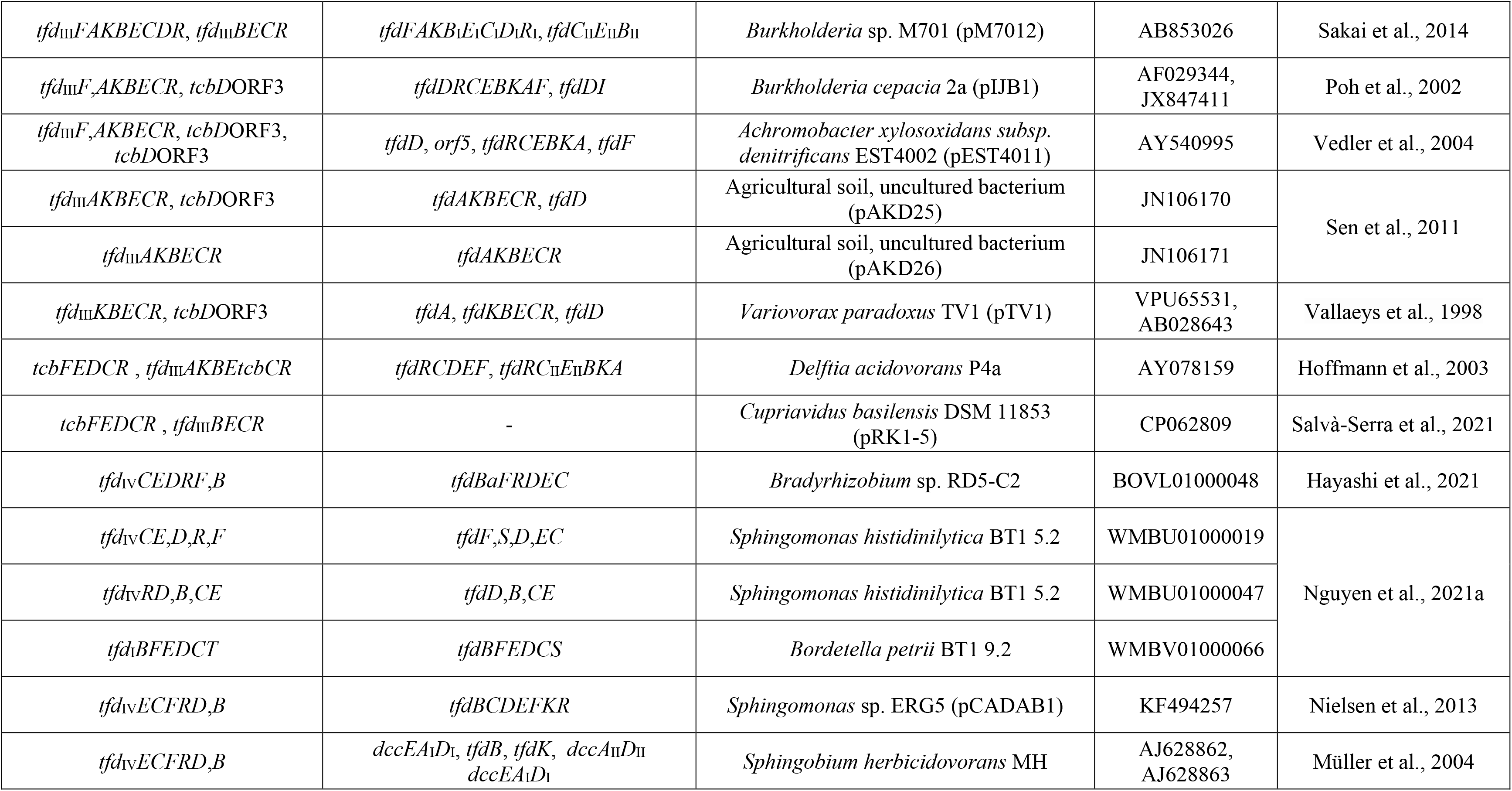
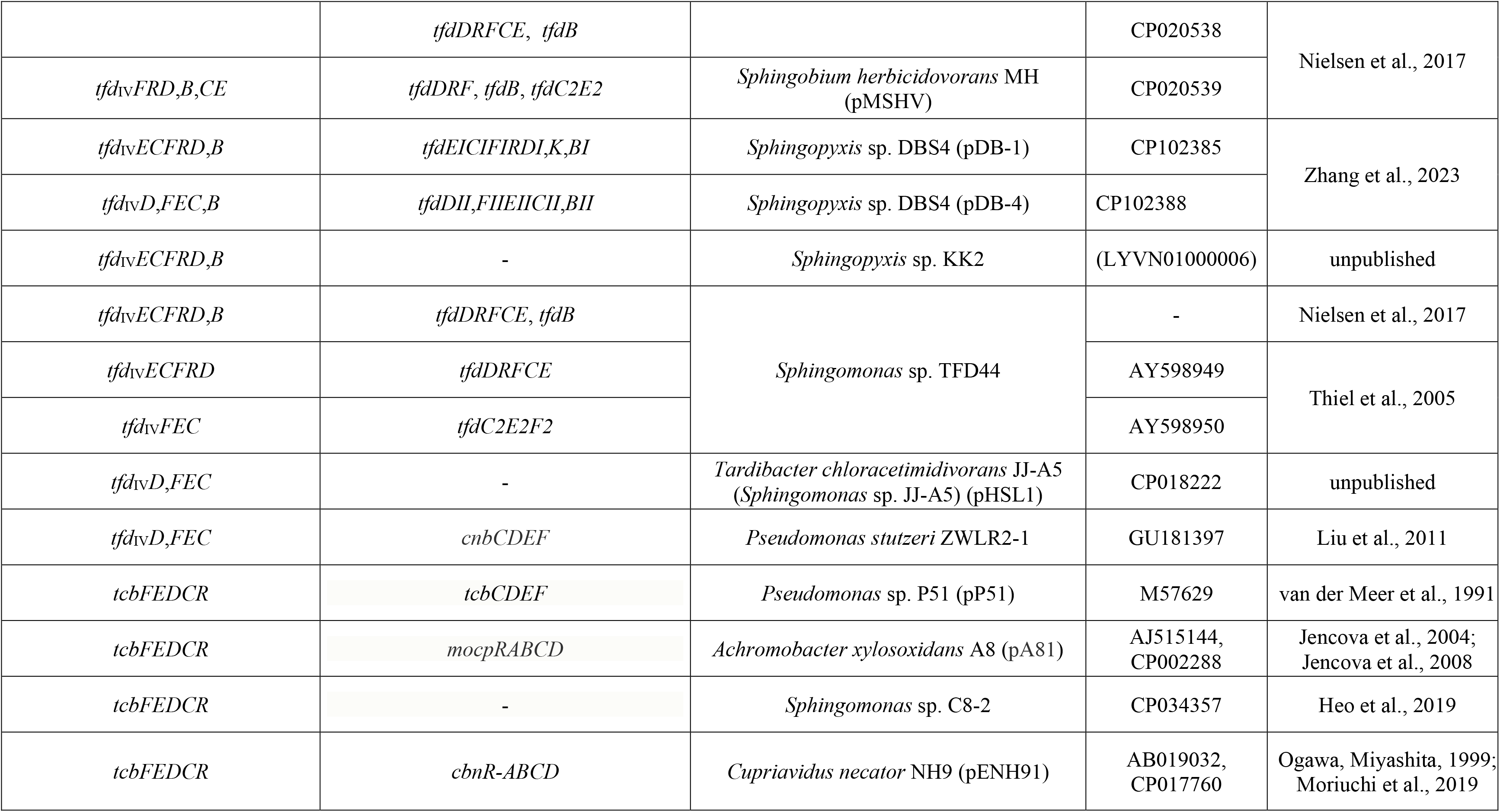
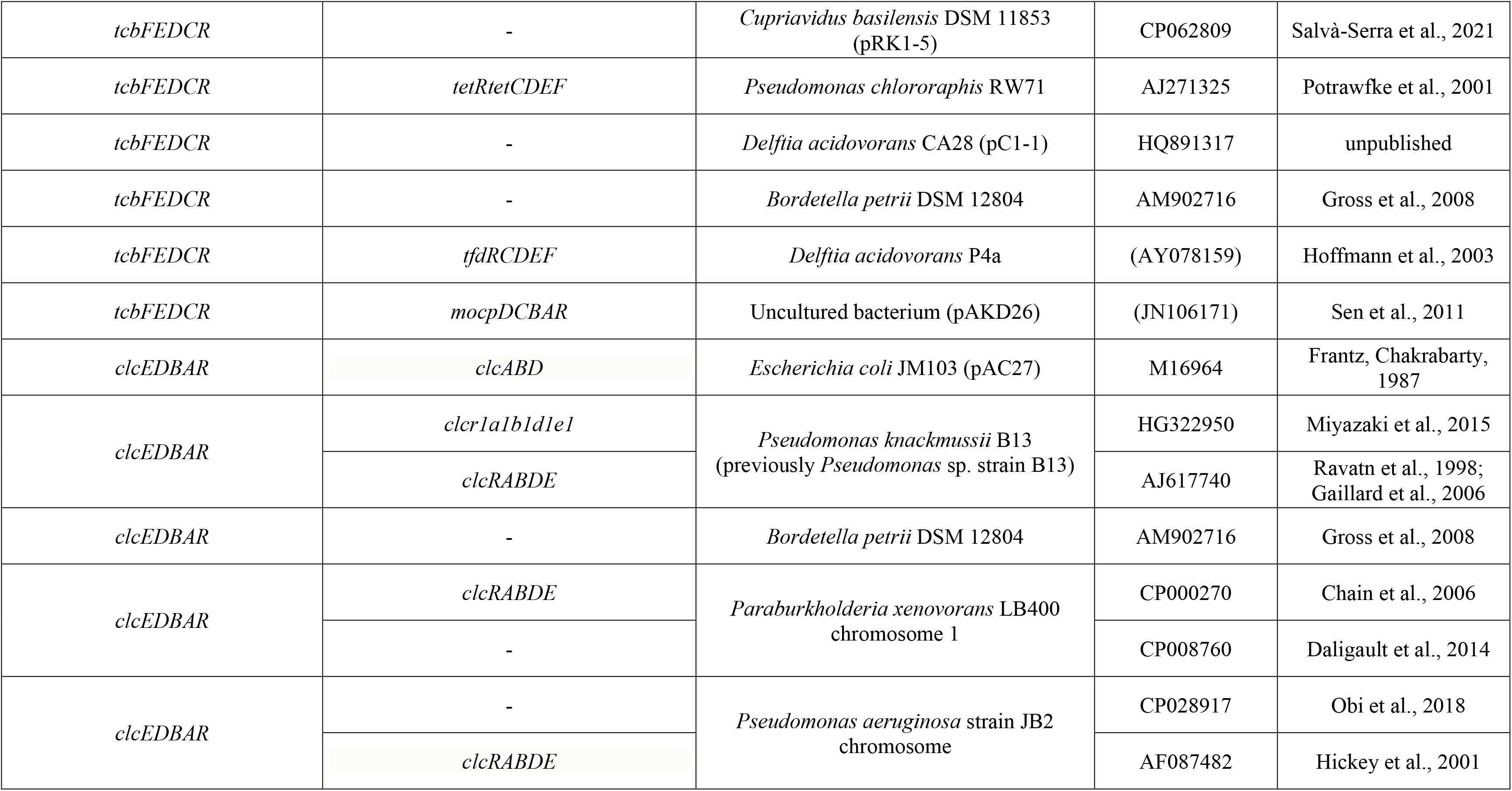
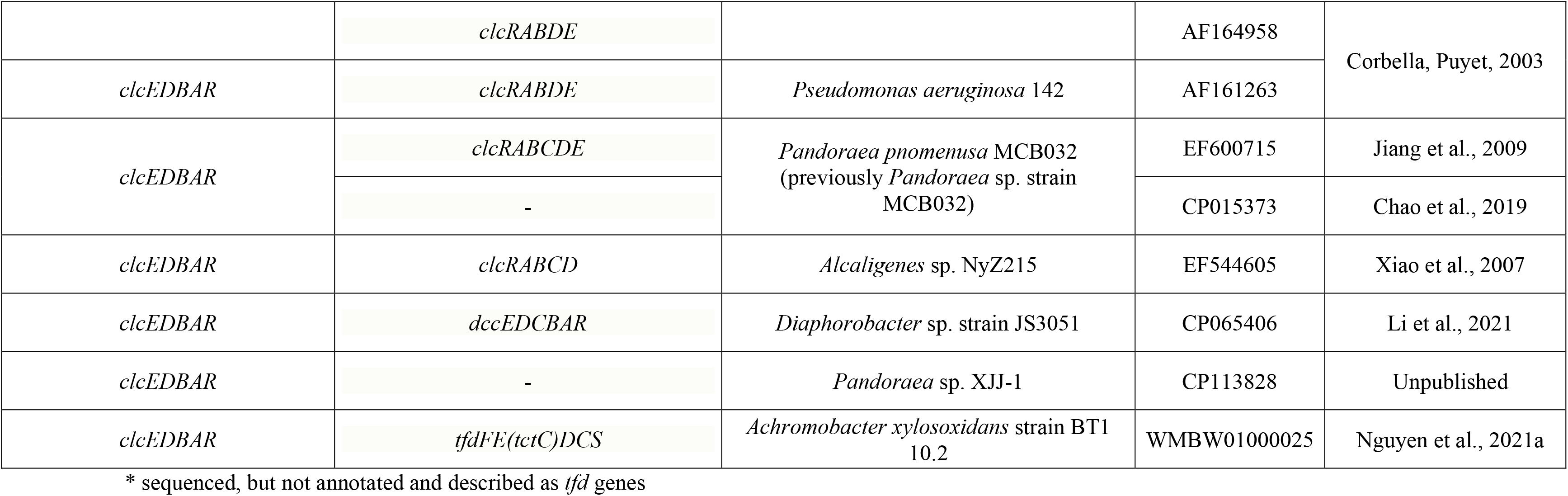
List of *tfd*, *tcb* and *clc* gene clusters.

The genetic structures of each *tfd*_I_ gene cluster were determined and are represented (fig. 1A). Although the genetic architecture *tfdB*_I_*F*_I_*E*_I_*D*_I_*C*_I_*T* is well conserved in most members of the *tfd*_I_ type, but subsequent comparative analyses suggested that another five clusters had an incomplete arrangement with deletions of the *tfdB*_I_ or *tfdT* genes. Amongst them were the *tfdF*_I_*E*_I_*D*_I_*C*_I_*T* cluster of the plasmid pNK84 (AP024328) and *P. phytofirmans* OLGA172 (*Burkholderia* sp. R172=*Burkholderia* sp. OLGA172) (CP014578). It should be noted that in *P. phytofirmans* OLGA172 earlier, an incomplete cluster with a *tfdD*_I_*C*_I_*TT* genetic architecture was found. A *tfdF*_I_*E*_I_*D*_I_*C*_I_ type structure was found in an uncultivated bacterium (AB478351). It is also interesting to note that in addition to the complete *tfd*_I_ gene cluster of the plasmid5, an incomplete cluster with the *tfdB*_I_*F*_I_*E*_I_*D*_I_ structure and fully identical at the DNA sequence level was found almost immediately downstream on the opposite strand of the plasmid (fig. 1A). Analysis of the DNA sequence identity of the aligned complete clusters of this type showed that it varied between 92.7% and 100%. At the same time, the pkk4 plasmid cluster had a minimum level of identity with respect to the other – the range was 92.7–96.8%. An interesting feature to note is that the *tfdT* gene, both in the pJP4 plasmid and in the vast majority of pPO line plasmids (with the exception of pPO4) was truncated. Thus, the vast majority of *tfd*_I_ type clusters are represented by a complete set of genes without genetic rearrangements.

**Fig. 1.**
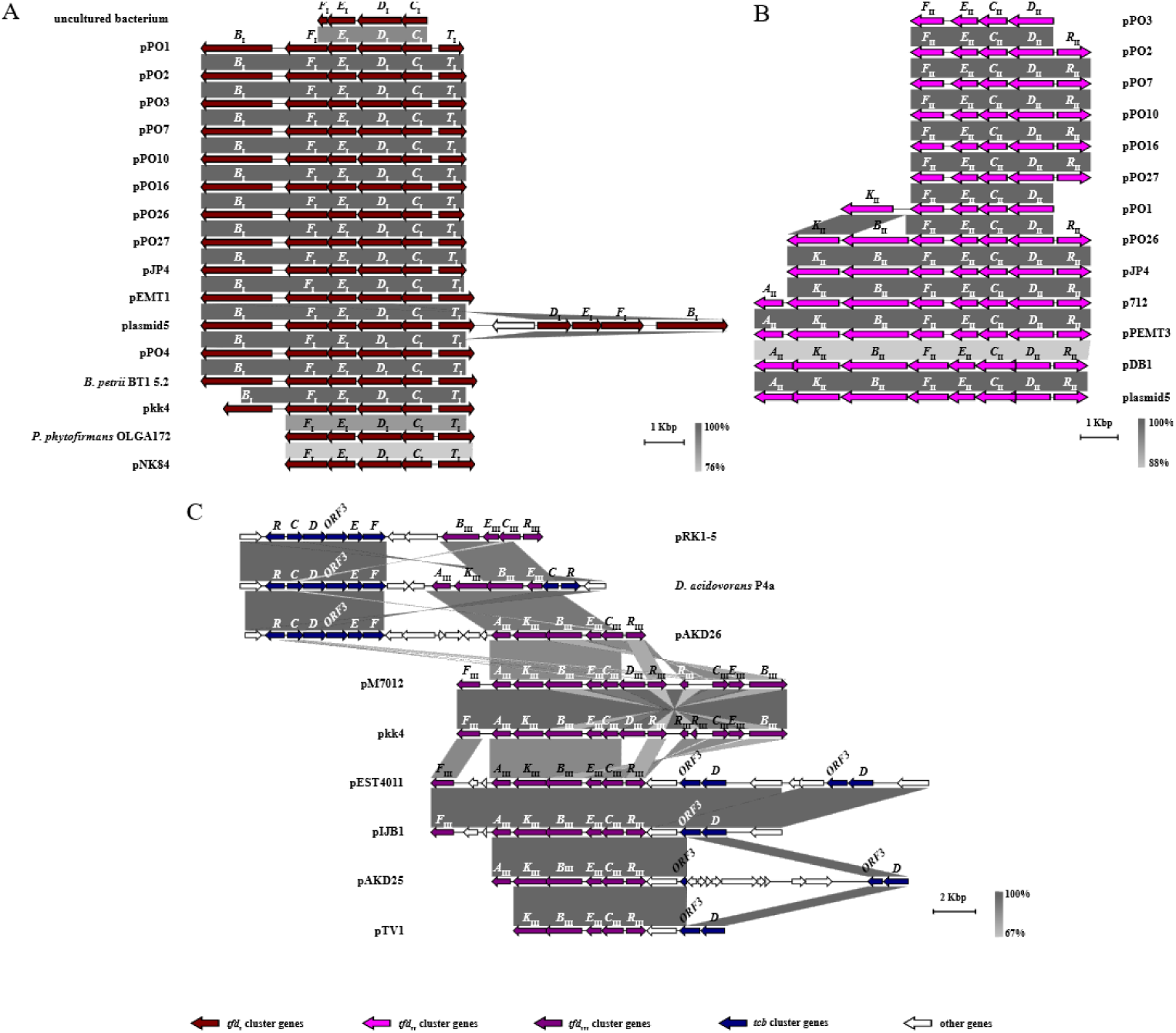
Comparative genomic analysis of (A) *tfd*_I_, (B) *tfd*_II_, and (C) *tfd*_III_ gene clusters showing their genomic rearrangements and evolutionary relationships between themselves and *tcb* gene clusters. Clusters and adjacent regions shown by linear visualization and open reading frames (ORFs) represented by arrows; the clusters are indicated by the color key (bottom). The degree of identity between clusters is indicated by the intensity of grayscale-shaded regions according to blastn as shown in the heat key (bottom right). The scale in kilobase pairs (Kbp) is shown at the bottom right of each cluster.

### The *tfd*_II_ gene cluster

A blastn search revealed thirteen clusters which had a genetic architecture closely related of the *tfd*_II_ seven-gene cluster of the pJP4 plasmid, namely *tfd*_II_*KBFECDR*. All the clusters had plasmid localization in bacterial hosts belonging to the *Burkholderiaceae* family. Four of them had a *tfd*_II_*AKBFECDR* eight-gene structure (plasmids pDB1 (JQ436721), p712 (JQ436722), pEMT3 (JX469827) and plasmid5 (CP038640)), with the *tfdA*_II_ gene located immediately downstream of the *tfdK*_II_ gene in a cluster, in contrast to the prototype plasmid pJP4. Thus, they had a complete set of genes in their structures (fig. 1B). It should be noted that the *tfdA*_II_ gene is absent in the *tfd*_II_ cluster in the pJP4 plasmid. It is located on the opposite strand downstream from *tfdR*_II_ gene of the cluster together with the *tfdS*_II_ gene and is separated from them by the open reading frames ORF31 and ORF32. However, unlike pJP4, three of them (pDB1, p712, and pEMT3) did not have the *tfd*_I_ gene cluster. Meanwhile, the plasmid5 plasmid contains the *tfd*_I_ cluster.

It should be pointed out that the *tfd*_II_*SA* clusters were also identified on the plasmid5, pkk4 (CP016005), pPO3 (CCJJ010000004), pPO4 (CCJK010000002), pPO10 (CCJM010000001) and pEMT1 plasmids which shared almost 100% identity at the nucleotide level. Separately, it is worth noting the variety of structures of the *tfd*_II_ type clusters on contigs for the pPO line plasmids (pPO1, 2, 3, 7, 10, 16, 26, 27). Of these, only pPO26 had the pJP4-like seven-gene cluster with the *tfd*_II_*KBFECDR* structure, while the others had structures genes: of the form *tfd*_II_*FECDR* (pPO2, 7, 10, 16, 27), *tfd*_II_*KFECD* (pPO1) and *tfd*_II_*FECD* (pPO3). A number of contigs had *tfd*_II_*SA* mini-clusters or *tfdA*_II_ and *tfdS*_II_ genes. Thus, twenty clusters with architectures related to *tfd*_II_*KBFECDR* and *tfd*_II_*SA* were identified. The six plasmids, namely pPO16, pJP4, p712, pEMT3, pDB1 and plasmid5, possessed core set of genes *tfd*_II_*KBFECDR* which shared identity values at the nucleotide level ranging from 87.9% to 100%.

### The *tfd*_III_ gene cluster

A large group of *tfd*_III_ type clusters (eleven) shared a common gene structure with different lengths has been identified by blastn-search (details of the new classification and nomenclature are described in the Materials and Methods and Discussion sections). Some of them were annotated without specifying the type of *tfd* cluster – *tfdDRCEBKAF* (Poh et al., 2002), *tfdRCDEF* (Hoffmann et al., 2003), *tfdRCEBKA* (Vedler et al., 2004), *tfdAKBECR* (Sen et al., 2011). Others belonged to the *tfd*_I_ and *tfd*_II_ type clusters **–** *tfdRC*_II_*E*_II_*BKA* (Hoffmann et al., 2003), *tfdFAKB*_I_*E*_I_*C*_I_*D*_I_*R*_I_, *tfdC*_II_*E*_II_*B*_II_ (Sakai et al., 2014) or were unpublished (plasmid pkk4) or not annotated (Salvà-Serra et al., 2021) (table 1). The more complete *tfd*_III_*FAKBECDR* eight-gene structures were encoded only on mega-plasmids pM7012 (AB853026) and pkk4. They were fully identical to each other and shared a 67.7– 68.2% identity with the complete clusters *tfd*_II_*AKBFECDR* for the *tfd*_II_ type (plasmids p712, pEMT3, pDB1 and plasmid5) at the nucleotide level. In both plasmids, the *tfdF*_III_ gene was located downstream of the *tfdA*_III_ gene. Another feature was the presence of incomplete versions of this type of cluster with structures *tfdBECRR* and *tfdBECR* immediately downstream of the *tfdR*_III_ gene on the opposite DNA strand. In these reduced clusters, the *tfdD*_III_ gene was completely deleted, while *tfdR*_III_ were partially reduced and, in pkk4, represented by two copies. The plasmid regions included both clusters on pM7012 and pkk4 almost identical at the nucleotide level. The clusters with structures *tfd*_III_*BECRR* and *tfd*_III_*BECR* both shared a 77.6% identity to with both *tfd*_III_*FAKBECDR* clusters. Importantly, in the mega-plasmid pkk4, the *tfd*_I_ type cluster was located upstream from *tfd*_III_*FAKBECDR*, as described below (fig. 1C).

The clusters with structures *tfd*_III_*F,AKBECR* were identified in plasmids pIJB1 (JX847411) and pEST4011 (AY540995). In them a *tfdF*_III_ genes were also located downstream of the *tfdA*_III_ gene like in the mega-plasmids pkk4 and pM7012 described above, but there were two small ORFs between them. Also, in contrast to pM7012 and pkk4, these clusters lacked a *tfdD*_III_ gene and short clusters *tfd*_III_*BECR*/*BECRR*. Both plasmids pAKD25 (JN106170) and pAKD26 (JN106171) from the same bacterial host (Sen et al., 2011) had clusters with a *tfd*_III_*AKBECR* gene structure without *tfdF*_III_ and *tfdD*_III_ genes. The shortest structures of that type were identified in plasmids pRK1-5 (CP062809), pTV1 (AB028643) and *Delftia acidovorans* P4a (AY078159). They had clusters with *tfd*_III_*KBECR*, *tfd*_III_*AKBE* and *tfd*_III_*BECR* gene structures, respectively. Interestingly, the core set of genes of this *tfd*_III_ cluster type (*tfd*_III_*BECR*), with the exception of *D. acidovorans* P4a, varied in terms of nucleotide identity by between 77.3% and 100%. An important distinguishing feature, in addition to a similar structure, in this group of clusters was the presence of *tcb* gene clusters adjacent to the *tfd*_III_ gene clusters. In total, nine clusters were identified, of which three had a full set of *tcbRCDEF* genes with ORF3, five clusters with structure *tcbD*ORF3 and one with *tcbRC*. The plasmids pkk4 and pM7012 were exceptions as they did not possess any *tcb* gene clusters. The complete *tcb* clusters were identified on opposite chains almost immediately downstream of the *tfdA*_III_ genes of *D. acidovorans* P4a and plasmid pAKD26, as well as the *tfdB*_III_ gene of plasmid pRK1-5. Moreover, in *D. acidovorans* P4a, the *tcbRC* cluster was identified immediately upstream of gene *tfdE*_III_. This was flanked by an ORF encoding a transcriptional regulator. The other plasmids: pEST4011, pIJB1, pAKD25, and pTV1 possessed a *tcbD*ORF3 cluster. Interestingly, pEST4011 possessed two copies of that cluster. The detailed synteny of the regions mentioned above is depicted in fig. 1C.

Moreover, identified in a number of plasmids (pEST4011, pIJB1, pAKD25 and pTV1) the two-gene clusters *tcbD*ORF3, were previously annotated as *tfdD* (Vallaeys et al., 1998), *tfdDI* (Poh et al., 2002), *tfdD* and orf5 (Vedler et al., 2004), *tfdD* and ORF (Sen et al., 2011). The analysis also revealed the presence of a unique two-gene *tcbCR* gene cluster adjacent to *tfd*_III_*AKBE* of *D. acidovorans* P4a previously annotated as *tfdRC*_II_ (Hoffmann et al., 2003).

### The *tfd*_IV_ gene clusters from *Sphingomonas* and *Bradyrhizobium*

This group combines non-traditional clusters that have variable gene structures and significantly differed from the canonical *tfd* gene clusters of the I and II types. The published clusters were annotated as *dccEA*_I_*D*_I_, *dccA*_II_*D*_II_, *dccEA*_I_*D*_I_ (Müller et al., 2004), *tfdC2E2, tfdC2E2F2*, *tfdDRFCE*, *tfdDRF* (Thiel et al., 2005; Nielsen et al., 2017), *cnbCDEF* (Liu et al., 2011), *tfdBCDEFKR* (Nielsen et al., 2013), *tfdEC*, *tfdCE* (Nguyen et al., 2021a), *tfdBaFRDEC* (Hayashi et al., 2021), *tfdEICIFIRDI*,*K*,*BI* and *tfdDII*,*FIIEIICII*,*BII* (Zhang et al., 2023). Nevertheless, comparative genomic analysis showed that the genetic structure of this *tfd*_IV_ type cluster could be divided into three main subtypes – A, B and C (fig. 2A) (details of the new classification and nomenclature are described in the Materials and Methods and Discussion sections). The most common structure, subtype A – *tfd*_IVA_*ECFRD*,*B*, possessed bacterial strains *Sphingobium herbicidovorans* MH, *Sphingopyxis* sp. KK2 and two plasmids pDB-1 and pCADAB1. According to Nielsen et al. (2017) the strain *Sphingomonas* sp. TFD44 also possesses *tfd*_IVA_*ECFRD*,*B*, but this sequence was absent in publicly available databases. Therefore, the early variant of the sequence *Sphingomonas* sp. TFD44 with a *tfd*_IVA_*ECFRD* structure without the *tfdB*_IVA_ gene which was sequenced by Thiel and colleagues (2005), was used for analysis. An interesting feature of *tfd*_IVA_*ECFRD*,*B* subclusters was the reverse orientation *tfdR*_IVA_ gene compared to the *tcb*, *clc* and other *tfd* clusters. Normally, that gene has the opposite orientation to other genes in the cluster. Moreover, the *tfdD*_IVA_ gene also had another orientation, opposite to the other cluster genes (with the exception of the pDB-1 plasmid). In the strain *Sphingomonas histidinilytica* BT1 5.2 (WMBU01000047) and on the plasmid pMSHV (CP020539) of the *S. herbicidovorans* MH, described above, less clustered variations with structures *tfd*_IVA_*RD*,*B*,*CE* and *tfd*_IVA_*FRD*,*B*,*CE*, respectively, were found. It should be noted that there is a high degree of synteny between the majority of this these subtype members (fig. 2B). The most common *tfd*_IVA_*ECFRD* structures of subtype A shared an identity at the nucleotide level ranging from 88.6% to 100%.

**Fig. 2.**
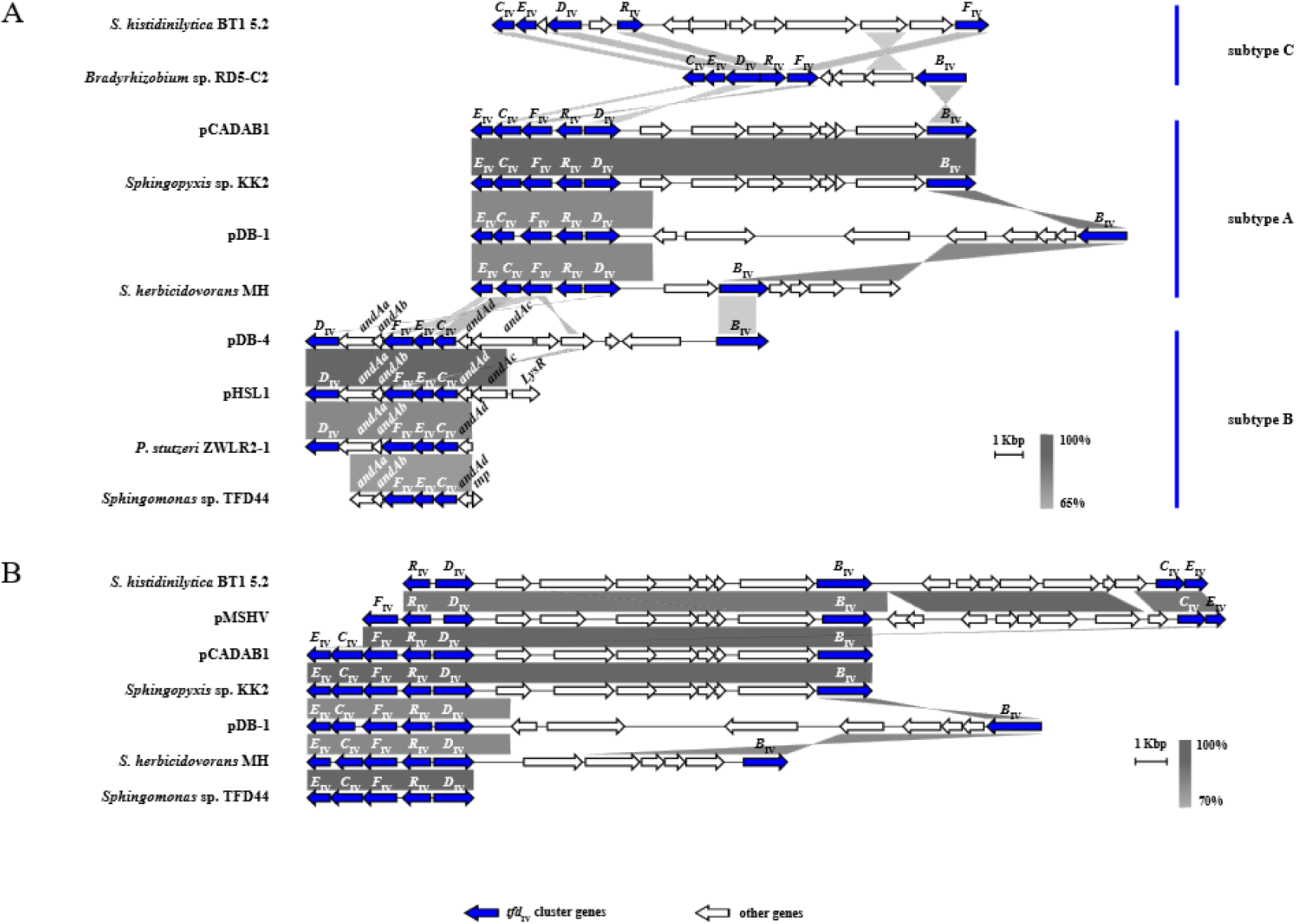
Comparative genomic analysis of *tfd*_IV_ gene clusters. (A) Genomic rearrangements and evolutionary relationships between subtypes A, B and C. (B) Synteny analysis showing putative gene assembly of *tfd*_IV_ gene cluster (subtype A). Clusters and adjacent regions shown by linear visualization and open reading frames (ORFs) represented by arrows; the clusters are indicated by the color key (bottom). The degree of identity between clusters is indicated by the intensity of grayscale-shaded regions according to blastn as shown in the heat key (bottom right). The scale in kilobase pairs (Kbp) is shown at the bottom right of each cluster.

The second subtype of this cluster, subtype B – *tfd*_IVB_*D*,*FEC/FEC*/*D*,*FEC*,*B*, was identified on the plasmids pDB-4 (CP102388), pHSL1 (CP018222) and in the strains *Pseudomonas stutzeri* ZWLR2-1 (GU181397) and *Sphingomonas* sp. tfd44. This last had the short variant without the *tfdD*_IVB_ gene – *tfd*_IVB_*FEC* of the whole set of genes in that subtype were ordered in one direction, in contrast to subtype A (fig. 2A). Interestingly, on the pHSL1 and pDB-4 gene structure, *tfd*_IVB_*FEC* was sandwiched into the cluster of genes responsible for encoding three-component Rieske-type [2Fe-2S] dioxygenase (anthranilate 1,2-dioxygenase) of *B. cepacia* DBO1 designated AntDO-3C and consisting of *AndAaAbAcAd* genes. The hybrid cluster has the structure *tfdD*_IVB_*AndAaAbtfd*_IVB_*FECAndAdAc*. Other strains, *P. stutzeri* ZWLR2-1 and *Sphingomonas* sp. tfd44, had shorter structures – *tfdD*_IVB_*AndAaAbtfd*_IVB_*FECAndAd* and *AndAaAbtfd*_IVB_*FECAndAd*, respectively. The *tfd*_IVB_*FEC* core set of genes shared 84.0–95.2% identity at the nucleotide level. Thus, that subtype was distributed mostly in the *Sphingomonadaceae* family with the exception of *P. stutzeri* ZWLR2-1.

The third subtype – C was presented on contigs of *Bradyrhizobium* sp. RD5-C2 (BOVL01000048) and *S. histidinilytica* BT1 5.2 (WMBU01000019) by gene structures with different degrees of gene assemblage – *tfd*_IVC_*CEDRF*,*B* and *tfd*_IVC_*CED*,*R*,*F*, respectively. The order and orientation of genes in that subtype differed from both subtype A and B (fig. 2A).

The obtained results indicated that three bacterial strains possessed two subtypes of *tfd*-like clusters. So, *Sphingomonas* sp. TFD44 (tfd44) and *Sphingopyxis* sp. DBS4 had the A and B subtypes, while *S. histidinilytica* BT1 5.2 had the A and C subtypes.

### The *tcb* and *clc* gene clusters

A blastn search revealed the sixteen sequences with high identity at nucleotide level with the canonical *clcRABDE* cluster of plasmid pP51 (M57629). Among them were both gene clusters annotated as *clc* and others: *clcr1a1b1d1e1* (Miyazaki et al., 2015), *dccEDCBAR* (Li et al., 2021) and *tfdFE(tctC)DCS* (Nguyen et al., 2021a). Additionally, some clusters were sequenced by different authors, but belonged to the same bacterial strain *Pseudomonas knackmussii* B13 (Ravatn et al., 1998; Gaillard et al., 2006; Miyazaki et al., 2015), *Paraburkholderia xenovorans* LB400 (Chain et al., 2006; Daligault et al., 2014), *Pseudomonas aeruginosa* strain JB2 (Hickey et al., 2001; Corbella, Puyet, 2003; Obi et al., 2018) and *Pandoraea pnomenusa* MCB032 (Jiang et al., 2009; Chao et al., 2019). As a result, the most fully sequenced gene clusters were taken for further analysis based on the latest data deposited in GenBank (table 1). From a genetic architecture point of view, all the clusters were highly conserved and had a common set of *clcRABDE* genes, as well as ORF3 genes in their structure. Within the *clc* cluster, the nucleotide sequences had a shared identity of between 96.3% and 100%. The plasmid pAC27 (M16964) *Pseudomonas aeruginosa* 142 (AF161263) possessed the truncated *clcR* gene. The ORF3 of strain *Diaphorobacter* sp. JS3051 was presented by the two truncated ORFs. There were no deletions, inversions or duplications of genes in the identified clusters. Fig. 3A illustrates the results of a comparative genomic analysis of the identified clusters and flanked ORFs. Moreover, the clusters of strains *B. petrii* DSM 12804 (AM902716), *P. knackmussii* B13 (HG322950), and plasmid unnamed 2 of the strain *P. pnomenusa* MCB032 (CP015373) had a high level of synteny downstream of the *clcE* gene. And downstream of the *clcR* gene, a high level of synteny was observed in *B. petrii* DSM 12804 (AM902716), P. *knackmussii* B13 (HG322950), *P. xenovorans* LB400 (CP008760), *P. aeruginosa* JB2 and *Diaphorobacter* sp. JS3051 (CP065406). Almost all the gene clusters, with one exception (pAC27, paaa and unnamed 2), had non-plasmid localization. Most of the bacteria carrying these clusters belonged to the families *Alcaligenaceae*, *Burkholderiaceae*, and *Comamonadaceae* of the order *Burkholderiales* of Betaproteobacteria, with the exception of few pseudomonades and *Escherichia coli* JM103 from Gammaproteobacteria.

**Fig. 3.**
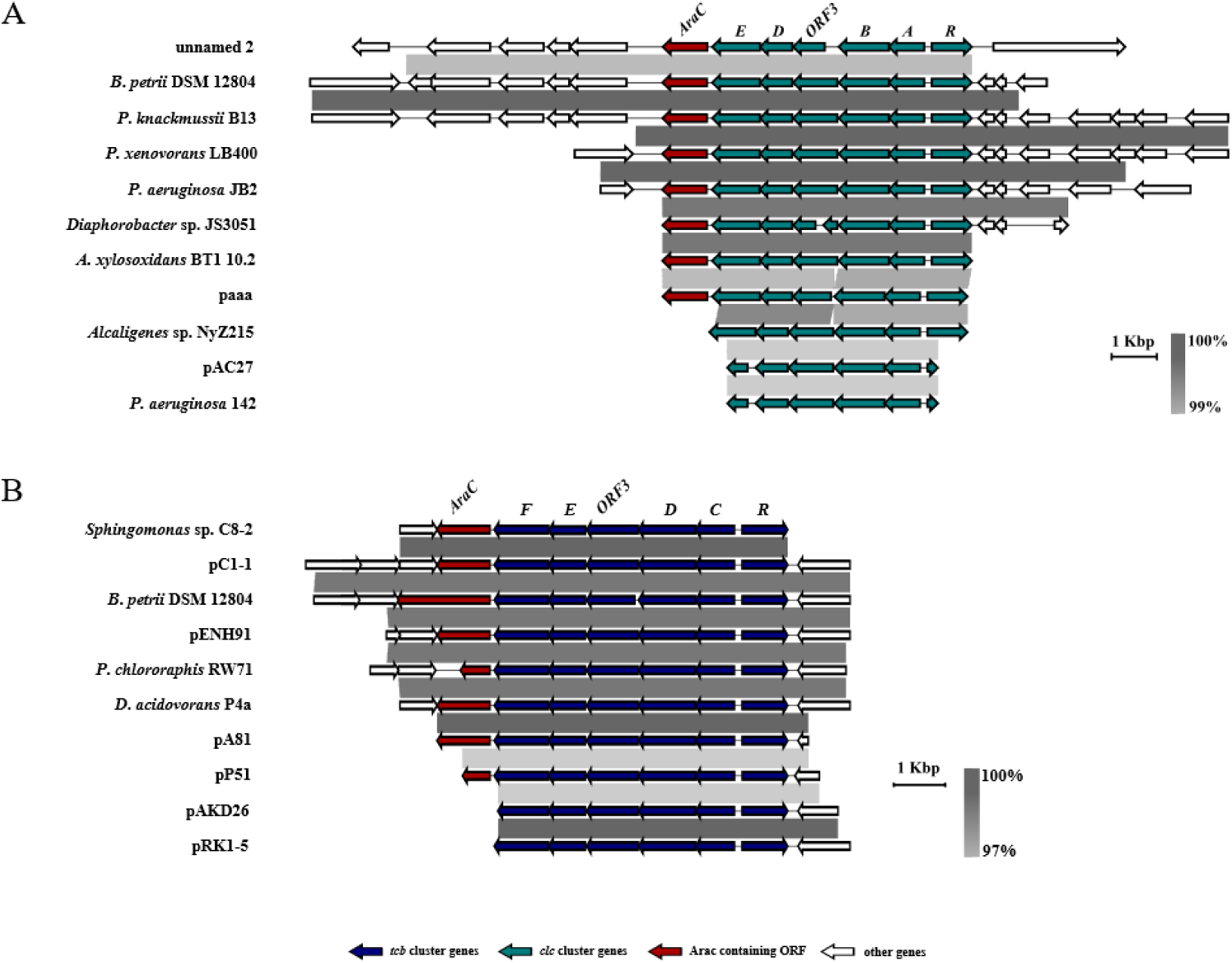
Comparative genomic analysis of (A) *clc* and (B) *tcb* gene clusters showing their synteny. Clusters and adjacent regions shown by linear visualization and open reading frames (ORFs) represented by arrows; the clusters are indicated by the color key (bottom). The degree of identity between clusters is indicated by the intensity of grayscale-shaded regions according to blastn as shown in the heat key (bottom right). The scale in kilobase pairs (Kbp) is shown at the bottom right of each cluster.

Additionally, blastn search indicated that the ten *tcb* clusters had a five-gene *RCDEF* structure (including the canonical *tcb* gene cluster of the plasmid pP51) which were mainly plasmid encoded: pP51 (M57629), pA81 (CP002288), pENH91 (CP017760), pRK1-5 (CP062809), pC1-1 (HQ891317), pAKD26 (JN106171). All the clusters shared between 97.0% and 100% identity and were much conserved in gene structure – they had no deletions, inversions or duplications. The four identified clusters were not annotated; the six clusters were designated as *tetRtetCDEF* (Potrawfke et al., 2001), *tfdRCDEF* (Hoffmann et al., 2003), *cbnR-ABCD* (Ogawa, Miyashita, 1999; Moriuchi et al., 2019), *mocpRABCD* (Jencova et al., 2008) and *mocpDCBAR* (Sen et al., 2011) (table 1).

Comparative genomic analysis has shown a high level of synteny of flanking ORFs for all the clusters (fig. 3B). Interestingly, only in genome *Bordetella petrii* DSM 12804 near the *RCDEF* was identified *tcbAB* gene cluster. Most of the bacteria carrying these clusters belonged to the families *Alcaligenaceae*, *Burkholderiaceae* and *Comamonadaceae* of the order *Burkholderiales* of Betaproteobacteria, with the exception of *Sphingomonas* sp. C8-2 и *Pseudomonas* sp. P51 from Alpha- and Gammaproteobacteria, respectively.

The common feature of both *clc* and *tcb* clusters was the flanking immediately downstream from the gene encoding maleylacetate reductases (*clcE* and *tcbF*, respectively) by a ORF with a contained conserved AraC binding domain. The following were the exceptions: strains *P. aeruginosa* 142, *Alcaligenes* sp. NyZ215 and plasmids pAC27, pAKD26 and pRK1-5. The ORFs across these clusters shared a similarity in the range of 42.5% to 100%. Meanwhile, the range of similarity across *clc* and *tcb* clusters ranged from 97.3% to 100% and from 71.5% to 100%, respectively.

### Comparative genomics for the *tfd*, *tcb* and *clc* clusters

Structurally, *clc*, *tcb* and all types of *tfd* clusters (excluding *tfd*_IV_) shared a common structure with a unidirectional set of genes, encoding catabolic reactions and regulatory protein in the opposite direction. Fig. 4 represents the comparative genomic analysis of the *clc* and *tcb* gene clusters of *B. petrii* DSM 12804, *tfd*_I_ cluster of plasmid5, *tfd*_II_ cluster of pEMT3, and *tfd*_III_ cluster of pkk4 plasmid. The *clc* and *tcb* clusters are closely related structurally. Both possess an ORF3 open reading frame with an unknown function and an identical order of genes encoding proteins with the same activity. Also, the *tfd*_I_ clusters are closely related to them, especially by tandem localization of genes encoding chloromuconate cycloisomerases (*clcD*, *tcbD* and *tfdD*_I_) and chlorocatechol 1,2-dioxygenases (*clcA*, *tcbC* and *tfdC*_I_). Nevertheless, there are two differences between *clc* and *tcb* and the *tfd*_I_ cluster namely in the presence of the *tfdB*_I_ gene and absence of ORF3. The results of multiple alignments of complete *clc*, *tcb* and *tfd*_I_ clusters revealed that *clc* and *tcb* shared between 46.2% and 50.2% similarity and between 47.5% and 50.2% similarity in terms of nucleotide level with *tfd*_I_, respectively. When comparing complete *clc* and *tcb* gene clusters, they were shown to have a similarity in the range of 62.0–63.2%.

**Fig. 4.**
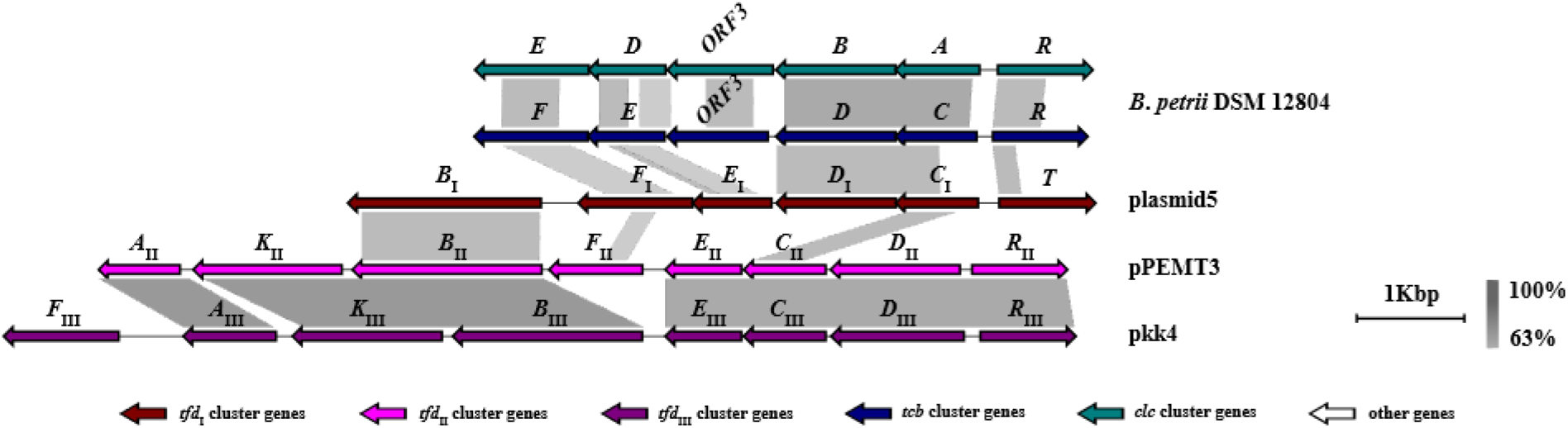
Comparison of gene structure of complete *tfd*, *clc* and *tcb* clusters. Clusters and adjacent regions shown by linear visualization and open reading frames (ORFs) represented by arrows; the clusters are indicated by the color key (bottom). The degree of identity between clusters is indicated by the intensity of grayscale-shaded regions according to blastn as shown in the heat key (bottom right). The scale in kilobase pairs (Kbp) is shown at the bottom right of each cluster.

Like *clc* and *tcb*, *tfd*_II_ and *tfd*_III_ clusters are structurally closely related to each other, but differ in terms of *tfdF* gene localization. Both *tfd*_II_ and *tfd*_III_ clusters possess a reverse order of chloromuconate cycloisomerase (*tfdD*_II_ and *tfdD*_III_) and chlorocatechol 1,2-dioxygenase (*tfdC*_II_ and *tfdC*_III_) encoding genes compared to *clc*, *tcb* and *tfd*_I_. But localization of the *tfdB* gene in *tfd*_I_, *tfd*_II_ and *tfd*_III_ clusters is identical. The similarity in terms of identity between *tfd*_II_ and *tfd*_III_ clusters ranged from 67.7% to 68.2%. At the same time *tfd*_I_ clusters shared from 49.0% to 52.2% and from 49.5% to 52.5% similarity in terms of identity at the nucleotide level with *tfd*_II_ and *tfd*_III_, respectively.

### The phylogenetic analysis of deduced protein sequences encoded by the *tfd*, *tcb* and *clc* gene clusters

Phylogenetic analysis of corresponding deduced protein sequences allowed for the evolutionary fate of genes in *tfd*, *tcb* and *clc* clusters, as well as the role of horizontal gene transfer to be assessed.

### *α*-ketoglutarate-dependent 2,4-D dioxygenase (*tfdA*)

Within α-ketoglutarate-dependent 2,4-D dioxygenases (TfdA) the ML analysis recovered *tfd*_II_ as a strongly-supported paraphyletic with *tfd*_III_ falling within that clade (fig. 5A). Interestingly, plasmid5 and pDB1 were recovered as sister clade with respect to plasmids in the *tfd*_II_ cluster with moderate support.

**Fig. 5.**
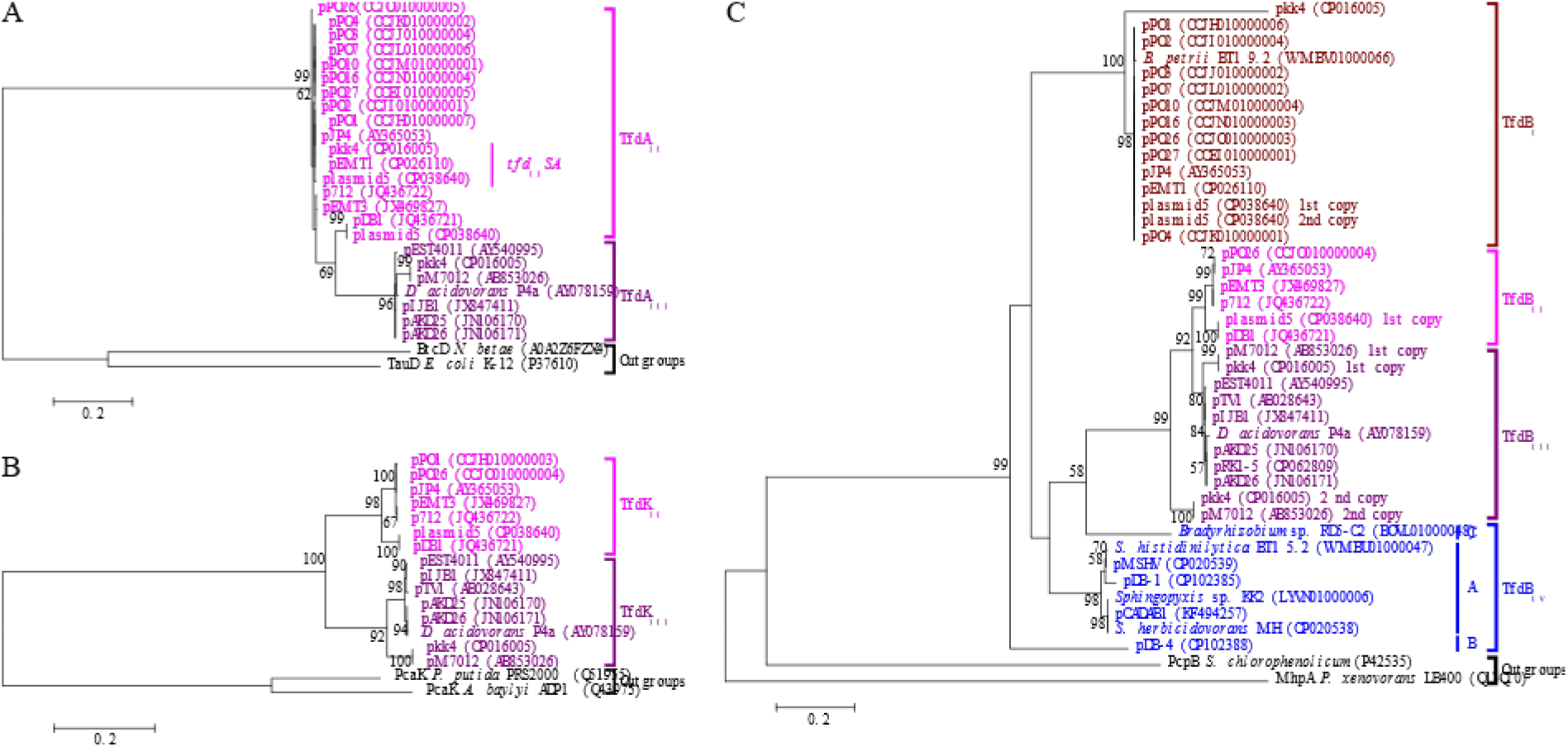
Phylogenetic classification of (A) alpha-ketoglutarate-dependent 2,4-dichlorophenoxyacetate dioxygenases (TfdA), (B) 2,4-D transport proteins (TfdK) and (C) 2,4-dichlorophenol hydroxylases (TfdB) of *tfd* gene clusters. Each color corresponds to one cluster. Bootstrap support from the maximum likelihood analyses (ML) higher than 50% are indicated above branches. The trees are drawn to scale, with branch lengths measured in the number of substitutions per site.

Comparative analysis of the protein sequences of α-ketoglutarate-dependent 2,4-D dioxygenases of the *tfd*_II_ and *tfd*_III_ clusters showed that similarity between them ranged from 87.2 % to 100%. The ranges of similarity between the protein sequences of TfdA’s inside each cluster were 93.8-100% and 96.9-100% for *tfd*_II_ and *tfd*_III_, respectively.

### 2,4-D transport protein (*tfdK*)

The ML tree topology recovered clades of *tfd*_II_ and *tfd*_III_ clusters as monophyletic with strong support. Each lineage further diverged into two well- and strongly-supported sister clades (fig. 5B). The two clades of the first lineage were formed TfdKs of plasmid5 and pDB1 on the one hand and p712, pEMT3, pJP4, pPO1 and pPO26 on the other. The second lineage diverged into two clades, the first of which was formed by almost all the transport proteins of the *tfd*_III_ clusters, and the second of which was formed by pkk4 and pM7012.

Comparative analysis of the amino acid sequences of 2,4-D transport proteins of the *tfd*_II_ and *tfd*_III_ clusters encoded by the *tfdK*_II_ and *tfdK*_III_ genes showed that similarity between them varied from 80.8% to 100%. The ranges of similarity between the amino acid sequences of TfdKs inside each cluster ranged from 94.4% to 100% and from 93.8% to 100% for *tfd*_II_ and *tfd*_III_ clusters, respectively.

### 2,4-DCP hydroxylase (*tfdB*)

The results of the ML analysis clearly showed strongly-supported paraphyly of the *tfd*_IV_ cluster (subcluster B) with respect to the *tfd*_I_, *tfd*_II_ and *tfd*_III_. Nevertheless, monophyly *tfd*_I_ clade and other clades were poorly supported (fig. 5C). Interestingly, subcluster A of the *tfd*_IV_ cluster resolved as paraphyletic with respect to the *tfd*_II_ and *tfd*_III_ clusters, but that result was not supported by bootstrap. Moreover, the *tfd*_III_ clade was recovered as a strongly-supported paraphyletic with respect to the *tfd*_II_ cluster.

A comparative analysis of amino acid sequences of 2,4-DCP hydroxylases encoded by the *tfdB*_I_, *tfdB*_II_, *tfdB*_III_, and *tfdB*_IV_ genes showed that they shared similarities of between 47.3% and 100%. The similarities between the amino acid sequences of TfdBs inside each cluster were in the range of 78.0–100% (*tfd*_I_), 95.9–100% (*tfd*_II_), 91.3–100% (*tfd*_III_), and 64.3– 100% (*tfd*_IV_), respectively.

### Chlorocatechol 1,2-dioxygenases (*tcbC*, *clcA*, *tfdC*_I_, *tfdC*_II_, *tfdC*_III_, *tfdC*_IV_)

The ML tree topology indicated that all the strongly-supported clades, uniting the chlorocatechol 1,2-dioxygenases of all the *tfd*, *clc* and *tcb* gene clusters (with the exception of subcluster C from the *tfd*_IV_ cluster) resolved as monophyletic with moderate support (fig. 6A). The clades uniting chlorocatechol 1,2-dioxygenases from the *tfd*_II_ and *tfd*_III_ clusters recovered as sister with strong support. The monophyly of chlorocatechol 1,2-dioxygenases for *tfd*_I_, *tfd*_II_ and *tfd*_III_ was not significantly supported. At the same time, clades uniting chlorocatechol 1,2-dioxygenases from the *clc* and *tcb* clusters resolved as moderately-supported sister clades. Within the *tfd*_III_ clade two well-supported sister clades were recovered, uniting proteins from plasmids pkk4 and pM7012 in the first clade and other plasmids in the second clade. The monophyly of clades, uniting proteins from subtypes A and B of the *tfd*_IV_ cluster was well supported.

**Fig. 6.**
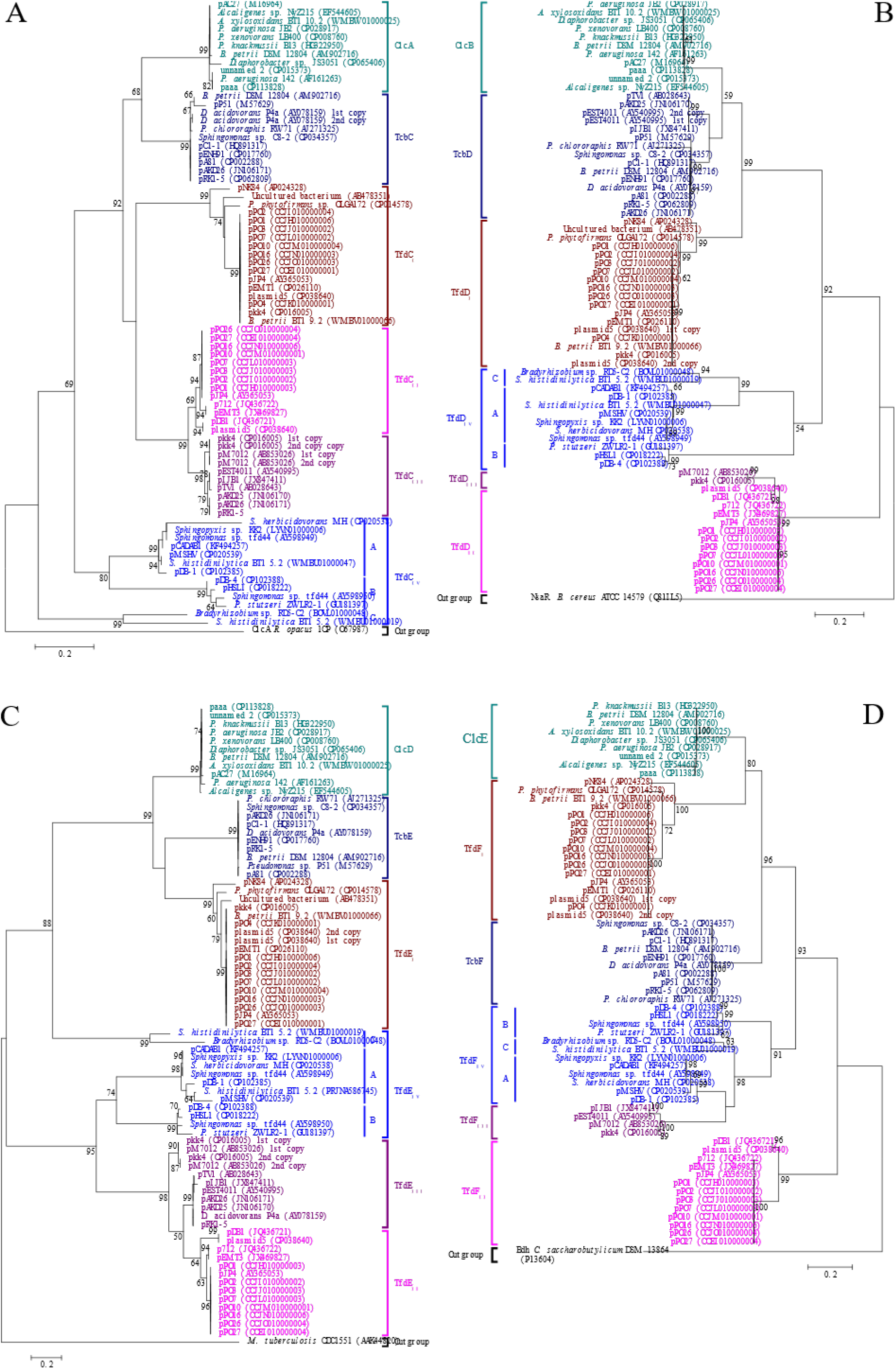
Phylogenetic classification of (A) chlorocatechol 1,2-dioxygenases, (B) chlormuconate cycloisomerases, (C) dienelactone hydrolases, (D) maleylacetate reductases of *tfd*, *tcb* and *clc* gene clusters. Each color corresponds to one cluster. Bootstrap support from the maximum likelihood analyses (ML) higher than 50% are indicated above branches. The trees are drawn to scale, with branch lengths measured in the number of substitutions per site.

A comparative analysis of the amino acid sequences of chlorocatechol 1,2-dioxygenases in these clusters encoded by the *tcbC*, *clcA*, *tfdC*_I_, *tfdC*_II_*, tfdC*_III_, and *tfdC*_IV_ genes showed that they shared a similarity of between 37.9% and 100%. The similarity between the amino acid sequences of chlorocatechol 1,2-dioxygenases inside each cluster was in the range of 95.2–100% (*tcb*), 96.9–100% (*clc*), 85.4–100% (*tfd*_I_), 95.7–100% (*tfd*_II_), 96.5–100% (*tfd*_III_) and 38.0–100% (*tfd*_IV_), respectively.

### Chloromuconate cycloisomerases (*tcbD*, *clcB*, *tfdD*_I_, *tfdD*_II_, *tfdD*_III_, *tfdD*_IV_)

In ML analysis, almost all the clades were resolved as well-supported monophyletic clades (fig. 6B). The exceptions to this were the two strongly-supported clades uniting the chloromuconate cycloisomerases from the *tfd*_II_ and *tfd*_III_ clusters. At the same time, *tfd*_II_ was recovered as a poorly supported paraphyletic clade with the *tfd*_III_ clade falling within her. In monophyletic lineage, all clades received strong support. At the same time, two sister clades with low support were recovered within the clade uniting chloromuconate cycloisomerases from the *tcb* cluster. The monophyly of the clades uniting chloromuconate cycloisomerases from *clc* and *tcb* was moderately supported. However, together with the *tfd*_I_ clade, they were recovered to be monophyletic with strong support. Within the *tfd*_IV_ cluster, three subclusters (A, B, and C), received good or strong support and were found to be monophyletic with moderate support.

A comparative analysis of the amino acid sequences of chloromuconate cycloisomerases in these clusters encoded by the *tcbD*, *clcB*, *tfdD*_I_, *tfdD*_II_, *tfdD*_III_, *tfdD*_IV_ genes indicated that they shared a similarity of between 45.6% and 100%. The similarity between the amino acid sequences of chloromuconate cycloisomerases inside each cluster was in the range of 80.3–100% (*tcb*), 98.7–100% (*clc*), 92.7–100% (*tfd*_I_), 94.8–100% (*tfd*_II_), 100% (*tfd*_III_) and 49.5–100% (*tfd*_IV_), respectively.

### Chlorodienelactone hydrolases (*tcbE*, *clcD*, *tfdE*_I_, *tfdE*_II_, *tfdE*_III_, *tfdE*_IV_)

The ML analysis of chlorodienelactone hydrolases were recovered as two monophyletic well- and strong-supported lineages (fig. 6C). The first included clusters *tcb*, *clc*, *tfd*_I_ and, surprisingly, subcluster A of the *tfd*_IV_ cluster. Each cluster formed a strongly-supported clade. The *tcb* and *tfd*_I_ clusters appear as the sister group, albeit that they received low support. The second lineage included *tfd*_II_, *tfd*_III_, and *tfd*_IV_ (subclusters A and B) clades with well- and strong support. The clade comprised *tfdE*_III_ chlorodienelactone hydrolases recovered as paraphyletic, with *tfd*_II_ cluster members falling within that clade. However, that result was moderately supported. Subclusters A and B of *tfd*_IV_ were recovered as sister clades with well-supported topology.

A comparative analysis of the amino acid sequences of chlorodienelactone hydrolases in these clusters encoded by the *tcbE*, *clcD*, *tfdE*_I_, *tfdE*_II_, *tfdE*_III_ and *tfdE*_IV_ genes indicated that they shared a similarity of between 25.5% and 100%. The similarity between the amino acid sequences of chlorodienelactone hydrolases inside each cluster varied from 99.6–100% (*tcb*), 97.0–100% (*clc*), 84.7–100% (*tfd*_I_), 88.1–100% (*tfd*_II_), 80.9–100% (*tfd*_III_) and 29.7–100% (*tfd*_IV_), respectively.

### Maleylacetate reductases (*tcbF*, *clcE*, *tfdF*_I_, *tfdF*_II_, *tfdF*_III_, *tfdF*_IV_)

This phylogenetic analysis recovered these proteins, encoded by all the analyzed clusters (with the exception of maleylacetate reductases from *tfd*_II_), as monophyletic with well support (Fig. 6D). As such, *clc*, *tcb*, *tfd*_I_, *tfd*_IV_ and *tfd*_III_, each clustered in well- and strongly-supported clades. Monophyly maleylacetate reductases from *clc*, *tcb* and *tfd*_I_ were strongly supported. The *tfd*_IV_ cluster was recovered as paraphyletic, with the *tfd*_III_ cluster falling within that clade albeit with low bootstrap support. The ML analysis recovered *tfd*_II_ as monophyletic with strong support and positioned the clade consisting of pDB1 and plasmid5 as the sister of the clade formed by all the other plasmids, also with strong support. All of the above confirmed the polyphyly of maleylacetate reductases.

A comparative analysis of the amino acid sequences of maleylacetate reductases in these clusters encoded by the *tcbF*, *clcE*, *tfdF*_I_, *tfdF*_II_, *tfdF*_III_, *tfdF*_IV_ genes indicated that they shared a similarity of between 49.7% and 100%. The ranges of similarity between the amino acid sequences of maleylacetate reductases inside each cluster were 98.8–100% (*tcb*), 89.5– 100% (*clc*), 84.8–100% (*tfd*_I_), 88.6–100% (*tfd*_II_), 96.9–100% (*tfd*_III_) and 60.7–100% (*tfd*_IV_).

### Transcriptional regulator (*tcbR*, *clcR*, *tfdT*, *tfdR*_II_/*tfdS*, *tfdR*_III_, *tfdR*_IV_)

The monophyly of the transcriptional regulators encoded by *clc*, *tcb*, *tfd*_I_, *tfd*_II_ and *tfd*_III_ clusters (with the exception of *tfd*_IV_) was well supported in ML analyses (fig. 7). The nodes, which are crucial for understanding the phylogenetic relationships between *clc*, *tcb*, *tfd*_I_, *tfd*_II_ and *tfd*_III_ clusters were weak or unsupported (with the exception of the *tfd*_II_ and *tfd*_III_ clusters). Nevertheless, each of the analyzed clusters, *tfd*_I_, *tfd*_II_, *tfd*_III_, *clc* and *tcb*, formed its own strongly-supported clade. The clade, consisting of TfdR_III_ transcriptional regulators, was recovered as a strongly-supported paraphyletic, with the *tfd*_II_ cluster members falling within that clade. The monophyly of *tfd*_IV_ cluster transcriptional regulators was moderately supported and within the *tfd*_IV_ cluster, both subclusters, I and III, were recovered to be monophyletic with strong and well support, respectively. Thus, transcriptional regulators were recovered as polyphyletic.

**Fig. 7.**
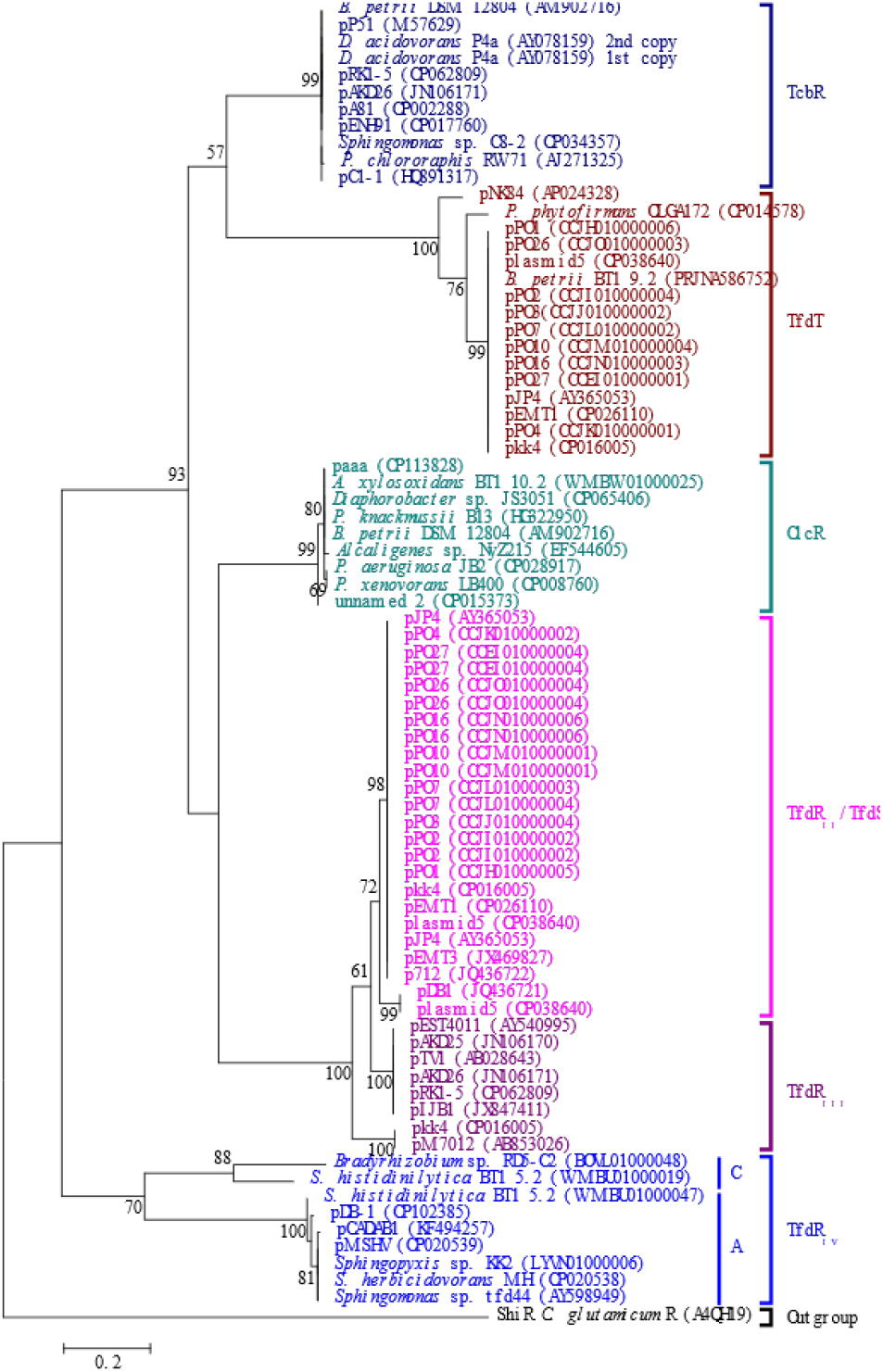
Phylogenetic classification of LysR-type transcriptional regulators (TfdR) of *tfd*, *tcb* and *clc* gene clusters. Each color corresponds to one cluster. Bootstrap support from the maximum likelihood analyses (ML) higher than 50% are indicated above branches. The trees are drawn to scale, with branch lengths measured in the number of substitutions per site.

A comparative analysis of the amino acid sequences of transcriptional regulators in these clusters encoded by the *tcbR*, *clcR*, *tfdT*, *tfdR*_II_/*tfdS*), *tfdR*_III_, *tfdR*_IV_ genes indicated that they shared a similarity of between 47.4% and 100%. The similarities between the amino acid sequences of maleylacetate reductases inside each cluster were in the ranges 99–100% (*tcb*), 96.6–100% (*clc*), 80.7–100% (*tfd*_I_), 95.6–100% (*tfd*_II_), 82.8–100% (*tfd*_III_) and 64.3–100% (*tfd*_IV_).

In summary, congruence between all the phylogenies of the proteins involved in catechol degradation through the ortho-cleavage pathway among the *tcb*, *clc* and *tfd*_I_ clusters can be concluded. The proteins of other clusters, *tfd*_II_, *tfd*_III_ and *tfd*_IV_, recovered both congruent and incongruent phylogenies. The proteins of cluster *tfd*_II_ were congruent with each other, as were the proteins of cluster *tfd*_III_, with one exception, namely, the protein maleylacetate reductase, which showed an incongruent phylogeny. Incongruence within cluster *tfd*_IV_ was observed among chlorocatechol 1,2-dioxygenases and chlorodienelactone hydrolase proteins.

## Discussion

### The new classification scheme and nomenclature of *tfd* gene clusters

The obtained results clearly indicated that *tfd*_I_ and *tfd*_II_ type clusters were well conserved in relation to their structures and could continue to be classified, as type I and type II, without any changes in classification. Meanwhile, the two groups of clusters are separate types of *tfd* clusters with substantive structural changes and different evolutionary origins. Taking this into account, they should be given independent designation numbers, type III and type IV. Thus, based on both the historical continuity of the designation of *tfd* gene clusters (Leveau et al., 1998; Laemmli et al., 2000) and the results of both comparative genomics and protein phylogeny, a new classification scheme is proposed, categorizing *tfd* gene clusters into four types – I, II, III and IV (A, B and C) alongside a new nomenclature (for details of the syntax, see the chapter Materials and Methods).

### The role of horizontal gene transfer (HGT) and gene displacement in the mosaic nature of *tfd* gene clusters

It is generally accepted that horizontal gene transfer (HGT) is the major process in bacterial evolution; its role has been proved in numerous studies. Mosaic operons contain genes transferred by HGT, which are characterized by the incongruence of their phylogeny with other genes (Omelchenko et al., 2003). Clusters, as well as operons, can also be mosaic in nature (Casjens, Thuman-Commike, 2011). Nevertheless, the analysis of *tfd* gene clusters from the standpoint of incongruence (or so-called discrepancies) in their phylogeny has only been performed in a few papers (McGowan et al., 1998; Vallaeys et al., 1999).

The congruence of the phylogeny of all *tcb* and *clc* cluster proteins clearly illustrates that they are not mosaic and are spread entirely by horizontal transfer. Moreover, the same conclusion can be drawn about the core five-gene part of cluster *tfd*_I_ responsible for the catechol ortho-cleavage pathway. Previously, it has been pointed out that from a phylogenetic point of view these clusters diverged from a common ancestral ortho-cleavage pathway for chlorocatechols (Schlömann, 1994).

Very intriguing findings follow from the congruence of almost all the proteins of clusters *tfd*_II_ and *tfd*_III_, with the exception of maleylacetate reductases (*tfdF*_II_ and *tfdF*_III_). These findings may be explained in the light of the HGT event, followed by differential gene losses – displacement of an ancestral *tfdF*_III_ (probably shared common ancestry with *tfdF*_II_) to the functionally equivalent gene of maleylacetate reductase, homologous to those from *tfdF*_IV_. This is a phenomenon, a homolog displacement, which is probably selectively neutral (Huang, Gogarten, 2008) but one of those that led to the origin of a different type of *tfd* cluster.

Analyzing the four proteins of cluster *tfd*_IV_ (TfdC_IV_, TfdD_IV_, TfdE_IV_ and TfdF_IV_) directly involved in the catechol pathway, it becomes obvious that subclusters A and B are not mosaic and are entirely distributed by HGT. In contrast, subcluster C, in the case of the chlorocatechol 1,2-dioxygenases (TfdC_IVC_) and chlorodienelactone hydrolases (TfdE_IVC_) proteins, had other evolutionary ancestors. These findings classify this subcluster as a mosaic.

Interestingly enough, the results of phylogenetic analysis clearly revealed that the *tfdB* gene for ancestral 2,4-DCP hydroxylase protein was assembled by clusters *tfd*_I_, *tfd*_II_, *tfd*_III_ and *tfd*_IV_ after distribution through HGT. However, with regard to gene *tfdB*, the results obtained in this work are inconsistent with an earlier conclusion about its independent evolution (Vallaeys et al., 1996).

Thus, HGT between orders *Burkholderiales* and *Sphingomonadales* provided phenomenal linkage between *tfd*_I_, *tfd*_II_, *tfd*_III_ and *tfd*_IV_ type clusters. And as a result, these orders have proven to be the most adapted to repeated exposures to the herbicide 2,4-D in soils.

### Evolution lineage including homologous *tfd*_I_, *tcb* and *clc* gene clusters

All the obtained results point to the existence of the lineage including three types of analyzed clusters, namely *tcb*, *clc* and *tfd*_I_ which are considered homologs. This conclusion is supported by at least five lines of evidence: (i) the clusters have similar core five-gene structures (genes, responsible for the ortho-cleavage of catechols) with an identical order of genes, especially encoded chloromuconate cycloisomerases and chlorocatechol 1,2-dioxygenases; (ii) the protein tree topologies suggest a common origin; (iii) high identities and similarities of DNA and protein sequences; (iv) the presence of ORF3 and the flanking ORF encoding the AraC family conserved domain in *tcb* and *clc*; (v) the clusters are widespread mainly across the order *Burkholderiales* (and even in a genome of the same strain, for example, *B. petrii* DSM 12804). Previous studies have suggested gene homology and possible evolutionary relatedness between genes of the first-described clusters *tcb*, *clc* and *tfd*_I_ (Ghosal, You, 1988; Coco et al., 1993; Kasberg et al., 1995).

The results apparently suggest that *tcb*, *clc* and *tfd*_I_ clusters are descendants of a single ancestral cluster, which further evolved by adapting to different substrates. The absence of ancestral cluster data, high conservation, and almost complete absence of genetic rearrangements among *tcb*, *clc* and *tfd*_I_ clusters prevent the causes and mechanisms of the main models of cluster formation from being delineated, with one exception. Nevertheless, the prevalent plasmid localization (with the exception of *clc* cluster) clearly suggests that they are the hot spots of cluster evolution and propagation. In turn, this indicates a possible assemblage of these clusters according to the ‘Scribbling Pad’ model (Norris, Merieau, 2013). The *clc* clusters are relatively poorly localized on plasmids compared to *tfd*_I_ and *tcb*, while at the same time remaining more conserved at the nucleotide sequence level. This also correlates well with this model. The *tcb* and *clc* gene clusters retained a high synteny of their own and flanking gene structure suggesting this is the ancestral state. Interestingly, the conservation of *tcb* cluster genes was higher than flanking ORFs that could suggest their high priority to bacterial hosts.

Perhaps *tfd*_I_ type clusters evolved from a common ancestor with clusters *tcb* and *clc* by clustering the *tfdB*_I_ gene (described above) and eliminating ORF3, resulting in the acquisition of the ability to hydroxylate 2,4-dichlorophenol (2,4-DCP) to 3,5-dichlorocatechol (3,5-DCC). This is supported by the absence of this gene in the clusters of the pNK84 plasmid and the *P. phytofirmans* OLGA172, which, according to the topology of all ortho-pathway proteins, diverged from a common ancestor earlier than the main group. Thus, they probably diverged before the *tfdB*_I_ gene was assembled. This allowed this cluster to more narrowly specialize in the degradation of 2,4-DCP into 3,5-DCC.

### Evolution lineage including homologous *tfd*_II_ and *tfd*_III_ type clusters

This lineage of *tfd* gene clusters is comprised of two types, *tfd*_II_ and *tfd*_III_, that have a similar structure and whose proteins are almost entirely homologous. The exception is the chloromaleylacetate reductase protein, which has a different evolutionary origin in those types (described above). There is no doubt that these types of clusters originated from the division of a common prototype cluster into two branches, as confirmed by the obtained results. By analogy with the evolutionary lineage including clusters *tcb*, *clc* and *tfd*_I_, only one of the potentially possible models of its formation can be distinguished for the prototype cluster, namely, the ‘Scribbling Pad’ model (Norris, Merieau, 2013), since, obviously, exclusively plasmid localization is a consequence of plasmid-mediated clusterization and propagation. Apparently, the crucial point in the evolutionary division of this lineage into two types was the homolog displacement of the *tfdF* gene in the common prototype cluster (described above).

Currently, it is obvious that the full *tfd*_II_ cluster with a complete *tfd*_II_*AKBFECDR* eight-gene structure is more widely distributed among plasmids than the *tfd*_II_*KBFECDR* seven-gene structure of the pJP4 plasmid. Nevertheless, the recent report about pPO line plasmids with the overall structure essentially the same as the pJP4 plasmid and with deletion variations of cluster *tfd*_II_ (Nguyen et al., 2019) puts a lot into perspective. Apparently, one consequent series of genetic rearrangements, primarily deletions, in pJP4 or pJP4-like plasmids, has resulted in the current diversity and propagation of these cluster variations. Since, of all the resulting variations, the seven-gene pJP4 cluster was the first to be explored, it became canonical.

The features typically associated with *tfd*_III_ type clusters can be derived through a series of rearrangements leading to incomplete clusters with *tfd*_III_*AKBECR* structure (excluding pRK1-5 and *D. acidovorans* P4a). It is probable that the ancient *tfd*_II_*AKBFECDR* exposed: (i) successful *tfdF*_II_ gene displacement that has resulted in a future *tfd*_III_*FAKBECDR* structure of the cluster (plasmids pkk4 and pM7012); (ii) simultaneously or sequentially successful displacement of the *tfdF*_II_ gene and unsuccessful displacement of *tfdD*_II_ that has resulted in a future *tfd*_III_*F*,*AKBECR* structure of the cluster (plasmids pIJB1 and pEST4011); (iii) simultaneously or sequentially unsuccessful displacement of the *tfdF*_II_ and *tfdD*_II_ gene that has resulted in a future *tfd*_III_*AKBECR* structure of the cluster (plasmids pAKD25, pAKD26, pTV1). It is important to note that plasmid pTV1 also has the *tfd*_III_*AKBECR* structure, but the *tfdA*_III_ gene was partially sequenced in further study (Vallaeys et al., 1996). These findings are not supported by the version proposed by Sakai et al. (2014) about the probability of the assemblage of the *tfd*_II_ type cluster recruiting genes from *tfd*_I_.

Moreover, the structure of *tfd*_III_*AKBECR* has undergone further rearrangements. In *D*. *acidovorans* P4a, the two-gene cluster *tfd*_III_*CR* was replaced with a two-gene cluster *tcbCR* which encoded the functionally identical proteins of the *tcb* cluster. This led to the emergence of a hybrid cluster *tfd*_III_*AKBEtcbCR*. Obviously, recombination has occurred between the *tfd*_III_ cluster and the *tcb* cluster located almost immediately downstream. Apparently, the plasmid pRK1-5 also had the structure *tfd*_III_*AKBECR*, but genes *tfd*_III_*AK* were deleted. Interestingly, plasmids pRK1-5, pAKD26 and *D. acidovorans* P4a have an almost similarly close proximity in terms of location to the *tfd*_III_ and *tcb* clusters, indicating that their ancestral state was *tcbRCDEF* and *tfd*_III_*AKBECR* and the other structure rearrangements occurred later. The latter is also indicated by the fact that plasmids pEST4011, pIJB1, pAKD25 and pTV1 have *tfd*_III_*AKBECR* and *tcb* clusters consisting of the ORF3 and *tcbD* gene instead of a complete *tcb* cluster. Finally, the ancestral state of *tcbRCDEF* and *tfd*_III_*AKBECR* is confirmed by the fact that plasmids pAKD25 and pAKD26 were isolated from the same bacterial host (Sen et al., 2011).

### Evolution lineage including unique *tfd*_IV_ type clusters

This *tfd*_IV_ clusters have a gene structure with incomplete clusterization, which significantly distinguishes them from the above-mentioned *tfd*_I_, *tfd*_III_, *clc* and *tcb* gene clusters. The second shared feature is several gene structures, and the absence of the *tfdA* gene in their structures. If the above clusters have a complete set of genes, then *tfd*_IV_ members have only five genes in the cluster. The results of both synteny and protein phylogenetic analysis revealed the three sublineages inside *tfd*_IV_ type clusters which evolved separately, but shared a common ancestor for the majority of encoded proteins.

The major subtype A involved seven clusters, three of which were plasmid-localized. The five of them shared five-gene structure and near localized *tfdB*_IVA_ gene *tfd*_IVA_*ECFRD*,*B* (fig. 2B) suggesting a common path for that sublineage clusterization. The induction of genes of that *tfd*_IV_ cluster sublineage in *Sphingomonas* sp. TFD44 in the presence of 2,4-D was previously noted by Thiel and colleagues (2005).

The members of the second subtype, B, of *tfd*_IV_ type clusters with the *tfd*_IVB_*D*,*FEC*/*FEC*/*D*,*FEC*,*B* structure for the cluster, to date, evolved as parts of pathways responsible for the degradation of multiple compounds. The presented results indicate that *tfd*_IVB_ type clusters are assembled with several genes encoding the anthranilate dioxygenase (AntDO-3C) enzyme of *B*. *cepacia* DBO1. Those genes are responsible for degradation of anthranilate (3-aminobenzoate) (Chang et al., 2003). Analogous recruitment was founded by Liu et al. (2011) for 2-chloronitrobenzene (2CNB) degrading genes. Nevertheless, the same cluster in *Sphingomonas* sp. TFD44 (*tfd*_IVB_*FEC* cluster) encoded 2,4-D degradation and had the *tfdD* gene in another place in the genome (Thiel et al., 2005). Since the above-mentioned clusters *tfd*_IVB_*D*,*FEC*/*FEC*/*D*,*FEC*,*B* had an almost identical structure, high identity and synteny, apparently, they distributed initially amongst order *Sphingomonadales* like other *tfd*_IV_ type clusters and later were acquired by *P*. *stutzeri* ZWLR2-1. The plasmid pDB-4 had the most complete *tfd*_IVB_*D*,*FEC*,*B* cluster and the presence of the *tfdB*_IV_ gene also indicated the relatedness of the first and second subtype of *tfd*_IV_ type clusters (fig. 2A). Additionally, it is probable that the above-mentioned dioxygenases showed broad substrate specificity. And *tfd*_IVB_*D*,*FEC*/*FEC*/*D*,*FEC*,*B* capable to degrading modified catechols which had come about as a result of the intermediate state of the catabolism of many compounds, including anthranilate, 2CNB, 2,4-D and others.

The third sublineage of *tfd*_IVC_ type clusters, *tfd*_IVC_*CEDRF*,*B/CE*,*D*,*R*,*F*, to date, can be defined by two clusters with different levels of clusterization in contigs of *Bradyrhizobium* sp. RD5-C2 (BOVL01000048) and *Sphingomonas histidinilytica* BT1 5.2 (WMBU01000019), respectively. The first one was designated as *tfdBaFRDEC* and involved in 2,4-D degradation from dichlorophenol with the same degradation pathway as that of *C*. *pinatubonensis* JMP134 (Hayashi et al., 2021). The *S*. *histidinilytica* BT1 5.2 which capable to degrade of 2,4-D possessed the second cluster, *tfd*_IVc_*CE*,*D*,*R*,*F*, which was annotated by Nguyen and colleagues (2021a) as *tfdF*,*S*,*D*,*EC*.

The main structural features, incomplete clusterization, across members of this lineage highlight the possible directions, driving forces and models of clusterization in contrast to other lineages. The extreme rarity of genes of this evolutionary lineage among bacteria may lead them to form clusters as a protective reaction in order to avoid evolutionary loss and promote propagation across bacteria. This would be very consistent with the ‘Selfish operon model’ (Lawrence, Roth, 1996) and apparently contradicts persistence as a driving force of their clustering (Fang et al., 2008). Moreover, the proven HGT, the absence of adjacent insertion elements (IS) with one exception and duplicated genes, actually oppose co-regulation theory (Price et al., 2005), the ‘IDE model’ (Kanai et al., 2022) and SNAP hypothesis (Brandis, Hughes, 2020), respectively. Interestingly, the revealed direct contribution of plasmids to genetic rearrangements in subtype A proves that a clusterization can proceed in two different models.

Currently, *Sphingomonas* and *Bradyrhizobium* genera are classified as I and III classes of 2,4-D degraders possessing the second system, *cadABCD* gene cluster, responsible for the initiation of degradation of chlorophenoxyacetic acids (Zharikova et al., 2018a). The *cad* clusters were identified in members of all *tfd*_IV_ subtypes. It was assumed that they were involved in the initial stages of 2,4-D degradation (Nielsen et al., 2013; Nielsen et al., 2017; Hayashi et al., 2021; Nguyen et al., 2021a). Nevertheless, some members of *Sphingomonas* and *Bradyrhizobium* could have their own versions of the first system, the *tfdA* gene and its homologs, named *tfdA*-like and *tfdAα*, respectively (Zharikova et al., 2018a).

Thus, *tfd*_IV_ clusters are unique *cad*-dependent 2,4-D degradation clusters which have evolved into three subtypes which provide *Sphingomonas* and *Bradyrhizobium* with great competitive advantages.

## Conclusion

The *tfd* is a unique family of diverse mosaic gene cluster types interlinked by their activity with regard to 2,4-D and other chlorinated aromatic compounds. Widespread mainly across the orders *Burkholderiales* and *Sphingomonadales*, the extraordinary reservoirs of genes involved in the ortho-cleavage pathway of 2,4-D and catechols, these diverse *tfds* as well as highly conserved *tcb* and *clc* gene clusters enable microbes to exploit a variety of xenobiotics-polluted niches. Systematization of both sequenced and published data for over forty years and subsequent analysis through comparative genomic and protein phylogeny approaches resulted in new insights in the evolution, classification and nomenclature of these clusters. Application of these work classification schemes provides a powerful approach for future exploration, especially for the correct annotation of *tfd*, *tcb* and *clc* clusters, as well as in the field related to their distribution and evolution across diverse bacteria.

## Funding

This work was supported by Russian Science Foundation (RSF) [grant number № 23-24-00480, https://rscf.ru/project/23-24-00480/].

## Notes

### Competing Interest Statement

The authors have declared no competing interest.

## References

Agency for Toxic Substances and Disease Registry (ATSDR). 2020. Toxicological profile for 2,4-Dichlorophenoxyacetic Acid (2,4-D). Atlanta, GA: U.S. Department of Health and Human Services, Public Health Service.

Brandis G, Hughes D. 2020. The SNAP hypothesis: Chromosomal rearrangements could emerge from positive Selection during Niche Adaptation. PLoS Genet. 16(3):e1008615. doi:10.1371/journal.pgen.1008615

Brucha G, Aldas-Vargas A, Ross Z, Peng P, Atashgahi S, Smidt H, Langenhoff A, Sutton NB. 2021. 2,4-Dichlorophenoxyacetic acid degradation in methanogenic mixed cultures obtained from Brazilian Amazonian soil samples. Biodegradation. 32(4):419–433. doi:10.1007/s10532-021-09940-3

Casjens SR, Thuman-Commike PA. 2011. Evolution of mosaically related tailed bacteriophage genomes seen through the lens of phage P22 virion assembly. Virology. 411(2):393–415. doi:10.1016/j.virol.2010.12.046

Chain PS, Denef VJ, Konstantinidis KT, Vergez LM, Agulló L, Reyes VL, Hauser L, Córdova M, Gómez L, González M. et al. 2006. *Burkholderia xenovorans* LB400 harbors a multi-replicon, 9.73-Mbp genome shaped for versatility. Proc Natl Acad Sci U S A. 103(42):15280–15287. doi:10.1073/pnas.0606924103

Chang HK, Mohseni P, Zylstra GJ. 2003. Characterization and regulation of the genes for a novel anthranilate 1,2-dioxygenase from *Burkholderia cepacia* DBO1. J Bacteriol. 185(19):5871–5881. doi:10.1128/JB.185.19.5871-5881.2003

Chao HJ, Chen YY, Wu J, Yan DZ, Zhou NY. 2019. Complete genome sequence of a chlorobenzene degrader, *Pandoraea pnomenusa* MCB032. Curr Microbiol. 76(11):1235–1237. doi:10.1007/s00284-019-01760-2

Coco WM, Rothmel RK, Henikoff S, Chakrabarty AM. 1993. Nucleotide sequence and initial functional characterization of the *clcR* gene encoding a LysR family activator of the *clcABD* chlorocatechol operon in *Pseudomonas putida*. J Bacteriol. 175(2):417–427. doi:10.1128/jb.175.2.417-427.1993

Corbella ME, Puyet A. 2003.Real-time reverse transcription-PCR analysis of expression of halobenzoate and salicylate catabolism-associated operons in two strains of *Pseudomonas aeruginosa*. Appl Environ Microbiol. 69(4):2269–2275. doi:10.1128/AEM.69.4.2269-2275.2003

Daligault HE, Davenport KW, Minogue TD, Bishop-Lilly KA, Broomall SM, Bruce DC, Chain PS, Coyne SR, Frey KG, Gibbons HS, et al. 2014. Whole-genome assemblies of 56 burkholderia species. Genome Announc. 2(6):e01106–14. doi: 10.1128/genomeA.01106-14.

Don RH, Weightman AJ, Knackmuss HJ, Timmis KN. 1985. Transposon mutagenesis and cloning analysis of the pathways for degradation of 2,4-dichlorophenoxyacetic acid and 3-chlorobenzoate in *Alcaligenes eutrophus* JMP134(pJP4). J Bacteriol. 161(1):85–90. doi:10.1128/jb.161.1.85-90.1985

Don RH, Pemberton JM. 1981. Properties of six pesticide degradation plasmids isolated from *Alcaligenes paradoxus* and *Alcaligenes eutrophus*. J Bacteriol. 145(2):681–686. doi:10.1128/jb.145.2.681-686.1981

Gaillard M, Vallaeys T, Vorhölter FJ, Minoia M, Werlen C, Sentchilo V, Pühler A, van der Meer JR. 2006. The *clc* element of *Pseudomonas* sp. strain B13, a genomic island with various catabolic properties. J Bacteriol. 188(5):1999–2013. doi: 10.1128/JB.188.5.1999-2013.2006.

Gross R, Guzman CA, Sebaihia M, dos Santos VA, Pieper DH, Koebnik R, Lechner M, Bartels D, Buhrmester J, Choudhuri JV, et al. 2008. The missing link: Bordetella petrii is endowed with both the metabolic versatility of environmental bacteria and virulence traits of pathogenic Bordetellae. BMC Genomics. 9:449. doi: 10.1186/1471-2164-9-449.

Fang G, Rocha EP, Danchin A. 2008. Persistence drives gene clustering in bacterial genomes. BMC Genomics. 9:4. doi:10.1186/1471-2164-9-4

Frantz B, Chakrabarty AM. 1987. Organization and nucleotide sequence determination of a gene cluster involved in 3-chlorocatechol degradation. Proc Natl Acad Sci U S A. 84(13):4460–4464. doi:10.1073/pnas.84.13.4460

Ghosal D, You IS. 1988. Nucleotide homology and organization of chlorocatechol oxidation genes of plasmids pJP4 and pAC27. Mol Gen Genet. 211(1):113–120. doi:10.1007/BF00338401

Ghosal D, You IS. 1989. Operon structure and nucleotide homology of the chlorocatechol oxidation genes of plasmids pJP4 and pAC27. Gene. 83(2):225–232. doi:10.1016/0378-1119(89)90108-x

Ghosal D, You IS, Chatterjee DK, Chakrabarty AM. 1985. Genes specifying degradation of 3-chlorobenzoic acid in plasmids pAC27 and pJP4. Proc Natl Acad Sci U S A. 82(6):1638–1642. doi:10.1073/pnas.82.6.1638

Harker AR, Olsen RH, Seidler RJ. 1989. Phenoxyacetic acid degradation by the 2,4-dichlorophenoxyacetic acid (TFD) pathway of plasmid pJP4: mapping and characterization of the TFD regulatory gene, *tfdR*. J Bacteriol. 171(1):314–320. doi:10.1128/jb.171.1.314-320.1989

Hayashi S, Tanaka S, Takao S, Kobayashi S, Suyama K, Itoh K. 2021. Multiple gene clusters and their role in the degradation of chlorophenoxyacetic acids in *Bradyrhizobium* sp. RD5-C2 isolated from non-contaminated soil. Microbes Environ. 36(3):ME21016. doi:10.1264/jsme2.ME21016

Heo J, Park I, You J, Han B-H, Kwon S-W, Lee S-W, Ahn J-H. 2019. Genome sequence analysis of *Sphingomonas histidinilytica* C8-2 degrading a fungicide difenoconazole. Korean J. Microbiol. 55(4):428–431. https://doi.org/10.7845/kjm.2019.9122

Hickey WJ, Sabat G, Yuroff AS, Arment AR, Pérez-Lesher J. 2001. Cloning, nucleotide sequencing, and functional analysis of a novel, mobile cluster of biodegradation genes from *Pseudomonas aeruginosa* strain JB2. Appl Environ Microbiol. 67(10):4603–4609. doi:10.1128/AEM.67.10.4603-4609.2001

Hoffmann D, Kleinsteuber S, Müller RH, Babel W. 2003. A transposon encoding the complete 2,4-dichlorophenoxyacetic acid degradation pathway in the alkalitolerant strain *Delftia acidovorans* P4a. Microbiology (Reading). 149(Pt 9):2545–2556. doi:10.1099/mic.0.26260-0

Huang J, Gogarten JP. 2008. Concerted gene recruitment in early plant evolution. Genome Biol. 9(7):R109. doi:10.1186/gb-2008-9-7-r109

Islam F., Wang J, Farooq MA, Khan MSS, Xu L, Zhu J, Zhao M, Muños S, Li QX, Zhou W. 2018. Potential impact of the herbicide 2,4-dichlorophenoxyacetic acid on human and ecosystems. Environ Int. 111:332–351. doi:10.1016/j.envint.2017.10.020

Jiang XW, Liu H, Xu Y, Wang SJ, Leak DJ, Zhou NY. 2009. Genetic and biochemical analyses of chlorobenzene degradation gene clusters in *Pandoraea* sp. strain MCB032. Arch Microbiol. 191(6):485–492. doi:10.1007/s00203-009-0476-9

Jencova V, Strnad H, Chodora Z, Ulbrich P, Hickey WJ, Paces V. 2004. Chlorocatechol catabolic enzymes from *Achromobacter xylosoxidans* A8. Internat Biodet Biodegrad 54, 175e181.https://doi.org/10.1016/j.ibiod.2004.03.007

Jencova V, Strnad H, Chodora Z, Ulbrich P, Vlcek C, Hickey WJ, Paces V. 2008. Nucleotide sequence, organization and characterization of the (halo)aromatic acid catabolic plasmid pA81 from *Achromobacter xylosoxidans* A8. Res Microbiol. 159(2):118–27. doi: 10.1016/j.resmic.2007.11.018.

Jones DT, Taylor WR, Thornton JM. 1992. The rapid generation of mutation data matrices from protein sequences. Comput Appl Biosci. 8(3):275–282. doi:10.1093/bioinformatics/8.3.275

Kanai Y, Tsuru S, Furusawa C. 2022. Experimental demonstration of operon formation catalyzed by insertion sequence. Nucleic Acids Res. 50(3):1673–1686. doi:10.1093/nar/gkac004

Kaphammer B, Kukor JJ, Olsen RH. 1990. Regulation of *tfdCDEF* by *tfdR* of the 2,4-dichlorophenoxyacetic acid degradation plasmid pJP4. J Bacteriol. 172(5):2280–2286. doi:10.1128/jb.172.5.2280-2286.1990

Kasberg T, Daubaras DL, Chakrabarty AM, Kinzelt D, Reineke W. 1995. Evidence that operons *tcb*, *tfd*, and *clc* encode maleylacetate reductase, the fourth enzyme of the modified ortho pathway. J Bacteriol. 177(13):3885–3889. doi:10.1128/jb.177.13.3885-3889.1995

Kim DU, Kim MS, Lim JS, Ka JO. 2013. Widespread occurrence of the *tfd-II* genes in soil bacteria revealed by nucleotide sequence analysis of 2,4-dichlorophenoxyacetic acid degradative plasmids pDB1 and p712. Plasmid. 69(3):243–248. doi:10.1016/j.plasmid.2013.01.003

Kumar A, Trefault N, Olaniran AO. 2016. Microbial degradation of 2,4-dichlorophenoxyacetic acid: Insight into the enzymes and catabolic genes involved, their regulation and biotechnological implications. Crit Rev Microbiol. 42(2):194–208. doi:10.3109/1040841X.2014.917068

Kumar S, Stecher G, Tamura K. 2016. MEGA7: Molecular Evolutionary Genetics Analysis version 7.0 for bigger datasets. Mol Biol Evol. 33(7):1870–1874. doi:10.1093/molbev/msw054

Laemmli CM, Leveau JH, Zehnder AJ, van der Meer JR. 2000. Characterization of a second *tfd* gene cluster for chlorophenol and chlorocatechol metabolism on plasmid pJP4 in *Ralstonia eutropha* JMP134 (pJP4). J Bacteriol. 182(15):4165–4172. doi:10.1128/JB.182.15.4165-4172.2000

Laemmli CM, Schönenberger R, Suter M, Zehnder AJ, van der Meer JR. 2002. TfdD_II_, one of the two chloromuconate cycloisomerases of *Ralstonia eutropha* JMP134 (pJP4), cannot efficiently convert 2-chloro-cis,cis-muconate to trans-dienelactone to allow growth on 3-chlorobenzoate. Arch Microbiol. 178(1):13–25. doi:10.1007/s00203-002-0417-3

Lawrence JG, Roth JR. 1996. Selfish operons: horizontal transfer may drive the evolution of gene clusters. Genetics. 143(4):1843–1860. doi:10.1093/genetics/143.4.1843

Leveau JH, Zehnder AJ, van der Meer JR. 1998. The *tfdK* gene product facilitates uptake of 2,4-dichlorophenoxyacetate by *Ralstonia eutropha* JMP134(pJP4). J Bacteriol. 180(8):2237–2243. doi:10.1128/JB.180.8.2237-2243.1998

Liu H, Wang SJ, Zhang JJ, Dai H, Tang H, Zhou NY. 2011. Patchwork assembly of nag-like nitroarene dioxygenase genes and the 3-chlorocatechol degradation cluster for evolution of the 2-chloronitrobenzene catabolism pathway in *Pseudomonas stutzeri* ZWLR2-1. Appl Environ Microbiol. 77(13):4547–4552. doi:10.1128/AEM.02543-10

Li T, Gao YZ, Xu J, Zhang ST, Guo Y, Spain JC, Zhou NY. 2021. A Recently assembled degradation pathway for 2,3-dichloronitrobenzene in *Diaphorobacter* sp. strain JS3051. mBio. 12(4):e0223121. doi: 10.1128/mBio.02231-21.

Liu S, Ogawa N, Miyashita K. 2001. The chlorocatechol degradative genes, *tfdT-CDEF*, of *Burkholderia* sp. strain NK8 are involved in chlorobenzoate degradation and induced by chlorobenzoates and chlorocatechols. Gene. 268(1-2):207–214. doi:10.1016/s0378-1119(01)00435-8

Madeira F, Pearce M, Tivey ARN, Basutkar P, Lee J, Edbali O, Madhusoodanan N, Kolesnikov A, Lopez R. 2022. Search and sequence analysis tools services from EMBL-EBI in 2022. Nucleic Acids Res. 50(W1):W276–W279. doi: 10.1093/nar/gkac240.

McGowan C, Fulthorpe R, Wright A, Tiedje JM. Evidence for interspecies gene transfer in the evolution of 2,4-dichlorophenoxyacetic acid degraders. Appl Environ Microbiol. 1998;64(10):4089–4092. doi:10.1128/AEM.64.10.4089-4092.1998

Miyazaki R, Bertelli C, Benaglio P, Canton J, De Coi N, Gharib WH, Gjoksi B, Goesmann A, Greub G, Harshman K, et al. 2015. Comparative genome analysis of *Pseudomonas knackmussii* B13, the first bacterium known to degrade chloroaromatic compounds. Environ Microbiol.17(1):91–104. doi: 10.1111/1462-2920.12498.

Morimoto S, Fujii T. 2009. A new approach to retrieve full lengths of functional genes from soil by PCR-DGGE and metagenome walking. Appl Microbiol Biotechnol. 83(2):389–396. doi:10.1007/s00253-009-1992-x

Moriuchi R, Dohra H, Kanesaki Y, Ogawa N. 2019. Complete genome sequence of 3-chlorobenzoate-degrading bacterium *Cupriavidus necator* NH9 and reclassification of the strains of the genera *Cupriavidus* and *Ralstonia* based on phylogenetic and whole-genome sequence analyses. Front Microbiol. 10:133. doi:10.3389/fmicb.2019.00133

Müller TA, Byrde SM, Werlen C, van der Meer JR, Kohler HP. 2004.Genetic analysis of phenoxyalkanoic acid degradation in *Sphingomonas herbicidovorans* MH. Appl Environ Microbiol. 70(10):6066–6075. doi:10.1128/AEM.70.10.6066-6075.2004

Nguyen TLA, Dang HTC, Koekkoek J, Braster M, Parsons JR, Brouwer A, de Boer T and van Spanning RJM. 2021. Species and metabolic pathways involved in bioremediation of vietnamese soil from Bien Hoa airbase contaminated with herbicides. Front. Sustain. Cities 3:692018. doi: 10.3389/frsc.2021.692018 (a)

Nguyen TLA, Dang HTC, Koekkoek J, Dat TTH, Braster M, Brandt BW, Parsons JR, Brouwer A and van Spanning RJM. 2021. Correlating biodegradation kinetics of 2,4-dichlorophenoxyacetic acid (2,4-D) and 2,4,5-trichlorophenoxyacetic acid (2,4,5-T) to the dynamics of microbial communities originating from soil in Vietnam contaminated with herbicides. Front. Sustain. Cities 3:692012. doi: 10.3389/frsc.2021.692012 (b)

Nguyen TPO, Hansen MA, Hansen LH, Horemans B, Sørensen SJ, De Mot R, Springael D. 2019. Intra- and inter-field diversity of 2,4-dichlorophenoxyacetic acid-degradative plasmids and their *tfd* catabolic genes in rice fields of the Mekong delta in Vietnam. FEMS Microbiol Ecol. 95(1). doi: 10.1093/femsec/fiy214

Nielsen TK, Rasmussen M, Demanèche S, Cecillon S, Vogel TM, Hansen LH. 2017. Evolution of sphingomonad gene clusters related to pesticide catabolism revealed by genome sequence and mobilomics of *Sphingobium herbicidovorans* MH. Genome Biol Evol. 9(9):2477–2490. doi:10.1093/gbe/evx185

Nielsen TK, Xu Z, Gözdereliler E, Aamand J, Hansen LH, Sørensen SR. 2013. Novel insight into the genetic context of the *cadAB* genes from a 4-chloro-2-methylphenoxyacetic acid-degrading *Sphingomonas*. PLoS One. 8(12):e83346. doi:10.1371/journal.pone.0083346

Norris V, Merieau A. 2013. Plasmids as scribbling pads for operon formation and propagation. Res Microbiol. 164(7):779–787. doi:10.1016/j.resmic.2013.04.003

Obi CC, Vayla S, de Gannes V, Berres ME, Walker J, Pavelec D, Hyman J, Hickey WJ. 2018. The integrative conjugative element *clc* (ICE*clc*) of *Pseudomonas aeruginosa* JB2. Front Microbiol. 9:1532. doi: 10.3389/fmicb.2018.01532.

Ogawa N, Miyashita K. 1999. The chlorocatechol-catabolic transposon Tn*5707* of Alcaligenes eutrophus NH9, carrying a gene cluster highly homologous to that in the 1,2,4-trichlorobenzene-degrading bacterium Pseudomonas sp. strain P51, confers the ability to grow on 3-chlorobenzoate. Appl Environ Microbiol. 65(2):724–731. doi:10.1128/AEM.65.2.724-731.1999

Omelchenko MV, Makarova KS, Wolf YI, Rogozin IB, Koonin EV. 2003. Evolution of mosaic operons by horizontal gene transfer and gene displacement in situ. Genome Biol. 4(9):R55. doi:10.1186/gb-2003-4-9-r55

Pérez-Pantoja D, Guzmán L, Manzano M, Pieper DH, González B. 2000. Role of *tfdC_I_D_I_E_I_F_I_* and *tfdD_II_C_II_E_II_F_II_* gene modules in catabolism of 3-chlorobenzoate by *Ralstonia eutropha* JMP134 (pJP4). Appl Environ Microbiol. 66(4):1602–1608. doi:10.1128/AEM.66.4.1602-1608.2000

Peterson MA, McMaster SA, Riechers DE, Skelton J, Stahlman PW. 2016. 2,4-D Past, Present, and Future: A Review. Weed Technol. 30(2):303–345. doi:10.1614/WT-D-15-00131.1

Plumeier I, Pérez-Pantoja D, Heim S, González B, Pieper DH. 2002. Importance of different *tfd* genes for degradation of chloroaromatics by *Ralstonia eutropha* JMP134. J Bacteriol. 184(15):4054–4064. doi:10.1128/JB.184.15.4054-4064.2002

Poh RP, Smith AR, Bruce IJ. 2002. Complete characterisation of Tn*5530* from *Burkholderia cepacia* strain 2a (pIJB1) and studies of 2,4-dichlorophenoxyacetate uptake by the organism. Plasmid. 48(1):1–12. doi:10.1016/s0147-619x(02)00018-5

Potrawfke T, Armengaud J, Wittich RM. 2001. Chlorocatechols substituted at positions 4 and 5 are substrates of the broad-spectrum chlorocatechol 1,2-dioxygenase of *Pseudomonas chlororaphis* RW71. J Bacteriol. 183(3):997–1011. doi:10.1128/JB.183.3.997-1011.2001

Pratama AA, Jiménez DJ, Chen Q, Bunk B, Spröer C, Overmann J, van Elsas JD. 2020. Delineation of a subgroup of the genus *Paraburkholderia*, including *P. terrae* DSM 17804^T^, *P. hospita* DSM 17164^T^, and four soil-isolated fungiphiles, reveals remarkable genomic and ecological features-proposal for the definition of a P. hospita species cluster. Genome Biol Evol. 12(4):325–344. doi: 10.1093/gbe/evaa031.

Price MN, Huang KH, Arkin AP, Alm EJ. 2005. Operon formation is driven by co-regulation and not by horizontal gene transfer. Genome Res. 15(6):809–819. doi:10.1101/gr.3368805

Ravatn R, Studer S, Zehnder AJ, van der Meer JR. 1998. Int-B13, an unusual site-specific recombinase of the bacteriophage P4 integrase family, is responsible for chromosomal insertion of the 105-kilobase *clc* element of *Pseudomonas* sp. strain B13. J Bacteriol. 180(21):5505–5514. doi:10.1128/JB.180.21.5505-5514.1998

Ricker N, Shen SY, Goordial J, Jin S, Fulthorpe RR. 2016. PacBio SMRT assembly of a complex multi-replicon genome reveals chlorocatechol degradative operon in a region of genome plasticity. Gene. 586(2):239–247. doi:10.1016/j.gene.2016.04.018

Sakai Y, Ogawa N, Shimomura Y, Fujii T. 2014. A 2,4-dichlorophenoxyacetic acid degradation plasmid pM7012 discloses distribution of an unclassified megaplasmid group across bacterial species. Microbiology (Reading). 160(Pt 3):525–536. doi:10.1099/mic.0.074369-0

Salvà-Serra F, Donoso RA, Cho KH, Yoo JA, Lee K, Yoon SH, Piñeiro-Iglesias B, Moore ERB, Pérez-Pantoja D. 2021. Complete multipartite genome sequence of the *Cupriavidus basilensis* type strain, a 2,6-dichlorophenol-degrading bacterium. Microbiol Resour Announc. 10(19):e00134–21. doi: 10.1128/MRA.00134-21.

Sayers EW, Bolton EE, Brister JR, Canese K, Chan J, Comeau DC, Farrell CM, Feldgarden M, Fine AM, Funk K, et al. 2023. Database resources of the National Center for Biotechnology Information in 2023. Nucleic Acids Res. 51(D1):D29–D38. doi: 10.1093/nar/gkac1032.

Sen D, Brown CJ, Top EM, Sullivan J. 2013. Inferring the evolutionary history of IncP-1 plasmids despite incongruence among backbone gene trees. Mol Biol Evol. 30(1):154–166. doi:10.1093/molbev/mss210

Sen D, Van der Auwera GA, Rogers LM, Thomas CM, Brown CJ, Top EM. 2011. Broad-host-range plasmids from agricultural soils have IncP-1 backbones with diverse accessory genes. Appl Environ Microbiol. 77(22):7975–7983. doi:10.1128/AEM.05439-11

Schlömann M. 1994. Evolution of chlorocatechol catabolic pathways. Conclusions to be drawn from comparisons of lactone hydrolases. Biodegradation. 5(3-4):301–321. https://doi.org/10.1007/BF00696467

Streber WR, Timmis KN, Zenk MH. 1987. Analysis, cloning, and high-level expression of 2,4-dichlorophenoxyacetate monooxygenase gene *tfdA* of *Alcaligenes eutrophus* JMP134. J Bacteriol. 169(7):2950–2955. doi:10.1128/jb.169.7.2950-2955.1987

Sullivan MJ, Petty NK, Beatson SA. 2011. Easyfig: a genome comparison visualizer. Bioinformatics. 27(7):1009–1010. doi:10.1093/bioinformatics/btr039

Thiel M, Kaschabek SR, Gröning J, Mau M, Schlömann M. 2005. Two unusual chlorocatechol catabolic gene clusters in *Sphingomonas* sp. TFD44. Arch Microbiol. 183(2):80–94. doi:10.1007/s00203-004-0748-3

Top EM, Holben WE, Forney LJ. 1995. Characterization of diverse 2,4-dichlorophenoxyacetic acid-degradative plasmids isolated from soil by complementation. Appl Environ Microbiol. 61(5):1691–1698. doi:10.1128/aem.61.5.1691-1698.1995

Trefault N, De la Iglesia R, Molina AM, Manzano M, Ledger T, Pérez-Pantoja D, Sánchez MA, Stuardo M, González B. 2004. Genetic organization of the catabolic plasmid pJP4 from *Ralstonia eutropha* JMP134 (pJP4) reveals mechanisms of adaptation to chloroaromatic pollutants and evolution of specialized chloroaromatic degradation pathways. Environ Microbiol. 6(7):655–668. doi:10.1111/j.1462-2920.2004.00596.x

van der Meer JR, Eggen RI, Zehnder AJ, de Vos WM. 1991. Sequence analysis of the *Pseudomonas* sp. strain P51 *tcb* gene cluster, which encodes metabolism of chlorinated catechols: evidence for specialization of catechol 1,2-dioxygenases for chlorinated substrates. J Bacteriol.173(8):2425–2434. doi:10.1128/jb.173.8.2425-2434.1991

van der Meer JR, Frijters AC, Leveau JH, Eggen RI, Zehnder AJ, de Vos WM. 1991. Characterization of the *Pseudomonas* sp. strain P51 gene *tcbR*, a LysR-type transcriptional activator of the *tcbCDEF* chlorocatechol oxidative operon, and analysis of the regulatory region. J Bacteriol. 173(12):3700–3708. doi:10.1128/jb.173.12.3700-3708.1991(b)

van der Meer JR, van Neerven AR, de Vries EJ, de Vos WM, Zehnder AJ. 1991. Cloning and characterization of plasmid-encoded genes for the degradation of 1,2-dichloro-, 1,4-dichloro-, and 1,2,4-trichlorobenzene of Pseudomonas sp. strain P51. J Bacteriol.173(1):6–15. doi:10.1128/jb.173.1.6-15.1991(c)

Vallaeys T, Albino L, Soulas G, Wright AD, Weightman AJ. 1998. Isolation and characterization of a stable 2,4-dichlorophenoxyacetic acid degrading bacterium, *Variovorax paradoxus*, using chemostat culture. Biotechnol Lett. 20:1073–1076 https://doi.org/10.1023/A:1005438930870

Vallaeys T, Courde L, Mc Gowan C, Wright AD, Fulthorpe RR. 1999. Phylogenetic analyses indicate independent recruitment of diverse gene cassettes during assemblage of the 2,4-D catabolic pathway. FEMS Microbiol Ecol. 28(4):373–382, https://doi.org/10.1111/j.1574-6941.1999.tb00591.x

Vallaeys T, Fulthorpe RR., Wright AM, Soulas G. 1996. The metabolic pathway of 2,4-dichlorophenoxyacetic acid degradation involves different families of *tfdA* and *tfdB* genes according to PCR-RFLP analysis. FEMS Microbiol Ecol. 20(3):163–172, https://doi.org/10.1111/j.1574-6941.1996.tb00315.x

Vedler E, Vahter M, Heinaru A. 2004. The completely sequenced plasmid pEST4011 contains a novel IncP1 backbone and a catabolic transposon harboring *tfd* genes for 2,4-dichlorophenoxyacetic acid degradation. J Bacteriol. 186(21):7161–7174. doi:10.1128/JB.186.21.7161-7174.2004

Xia ZY, Zhang L, Zhao Y, Yan X, Li SP, Gu T, Jiang JD. 2017. Biodegradation of the herbicide 2,4-dichlorophenoxyacetic acid by a new isolated strain of *Achromobacter* sp. LZ35. Curr Microbiol. 74(2):193–202. doi: 10.1007/s00284-016-1173-y.

Xiang S, Lin R, Shang H, Xu Y, Zhang Z, Wu X, Zong F. 2020. Efficient degradation of phenoxyalkanoic acid herbicides by the alkali-tolerant *Cupriavidus oxalaticus* strain X32. J Agric Food Chem. 68(12):3786–3795. doi: 10.1021/acs.jafc.9b05061

Xiao Y, Zhang JJ, Liu H, Zhou NY. 2007. Molecular characterization of a novel ortho-nitrophenol catabolic gene cluster in *Alcaligenes* sp. strain NyZ215. J Bacteriol. 189(18):6587–6593. doi:10.1128/JB.00654-07

Yamamoto-Tamura K, Moriuchi R, Ogawa N. 2021.Complete genome sequence of *Caballeronia* sp. strain NK8 (MAFF311271), a chlorobenzoate-degrading bacterium. Microbiol Resour Announc. 10(31):e0041621. doi:10.1128/MRA.00416-21

Zhang L, Song M, Mao Z, Liu Y, Li F, Jiang J, Chen K. 2023. A new enantioselective dioxygenase for the (S)-enantiomer of the chiral herbicide dichlorprop in *Sphingopyxis* sp. DBS4. Int Biodeterior Biodegradation, 176:105511, https://doi.org/10.1016/j.ibiod.2022.105511.

Zharikova NV, Iasakov TR, Zhurenko EI, Korobov VV, Markusheva TV. 2018. Bacterial genes of non-heme iron oxygenases, which have a Rieske-type cluster, catalyzing initial stages of degradation of chlorophenoxyacetic acids. Russ J Genet. 54(3):284–295. doi: 10.1134/S1022795418030171 (a)

Zharikova NV, Iasakov TR, Zhurenko EI, Korobov VV, Markusheva TV. 2021. Plasmids of the chlorophenoxyacetic-acid degradation of bacteria of the genus *Raoultella*. Appl Biochem Microbiol. 57:335–343. https://doi.org/10.1134/S0003683821030157

Zharikova NV, Iasakov TR, Zhurenko EYu, Korobov VV, Markusheva TV. 2018. Bacterial genes of 2,4-dichlorophenoxyacetic acid degradation encoding α-ketoglutarate-dependent dioxygenase activity. Biol Bull Rev8(2):155–167. doi: 10.1134/S2079086418020081 (b)

